# Positive and relaxed selective pressures have both strongly influenced the evolution of cryonotothenioid fishes during their radiation in the freezing Southern Ocean

**DOI:** 10.1101/2022.02.01.478646

**Authors:** Kevin T. Bilyk, Xuan Zhuang, Chiara Papetti

## Abstract

Evolution in the chronic cold of the Southern Ocean has had a profound influence on the physiology of cryonotothenioid fishes. However, the suite of genetic changes underlying the physiological gains and losses in these fishes is still poorly surveyed. Using molecular evolution techniques, this study aimed to identify which functional classes of genes changed during the cryonotothenioid radiation in a polar ocean. The influences of both positive and relaxed selective pressures were isolated following two major physiological transitions: the onset of freezing temperatures, and the loss of hemoproteins. Looking at the influence of cold temperatures, positive selective pressure was most prominently found to act on biosynthetic processes (the machinery of transcription and translation) as well as on protein polymerization, cell adhesion, and cell cycle progression, suggesting these are key challenges to life in freezing waters. Relaxation of selective pressure produced a more complex pattern of change, affecting several homeostatic processes, suggesting their attenuation in cold-stable and oxygen-rich waters, but also seemingly acting as a route to key genetic change behind the morphological and ecological diversification seen in the clade. Finally, while chronic cold-water temperatures appear to have instigated clear genetic change, the loss of hemoproteins led to little observable change relative to their red-blooded relatives. Combined, the influence of positive and relaxed selection show that long-term exposure to cold has led to profound changes in cryonotothenioid genomes, which may make it challenging for them to respond to unpredictable heat waves and to adapt to a rapidly changing climate.

## INTRODUCTION

Today, the waters of the Antarctic shelf are dominated by the members of a single taxonomic group, the cryonotothenioid fishes comprised of the five Antarctic families of the perciform suborder Notothenioidei (Dornburg et al, 2017; Eastman, 2005; Near et al, 2018). The origin of this group is closely linked to the onset of freezing conditions in Antarctic waters (Matschiner et al, 2011), and their subsequent radiation driven by continued changes to the region’s climate (Near et al, 2012). Evolution in this isolated, frigid environment has resulted in a diverse group of fishes that now share a profound specialization to life in the cold (Beers and Jayasundara, 2015). However, the shared genetic consequences of evolution in chronic cold remain poorly surveyed in this group, as are the genetic changes that may have come as the cryonotothenioids diversified in the Southern Ocean.

The most obvious change in selective pressure on the cryonotothenioids comes from the temperature of the Southern Ocean. Surface water temperatures remain below 1.5°C, and freezing water temperatures define the most species rich waters along the Antarctic coast, even to great depths (DeVries and Steffensen, 2005). In addition to being cold, the waters around Antarctica are remarkable for the stability of these cold temperatures, with only modest thermal variability compared to temperate and tropical waters (Barnes et al, 2006). This reaches an extreme in high-latitude habitats, such as the waters of McMurdo Sound, which are characterized by water temperatures that remain near their freezing point throughout the year (Cziko et al, 2014).

Although the origin of the cryonotothenioid clade is estimated at 22.4 Ma (Near et al, 2012), most of the living diversity of Antarctic notothenioids has originated more recently, following the onset of the Middle Miocene climatic transition (MMCT) that initiated modern subzero polar conditions between 14.2 and 13.8 Ma (Shevenell et al, 2004). Evolution in these freezing waters would expectantly have imposed strong selective pressures to deal with the biological challenges of life in the cold, but it would also have exposed endemic fishes to an important relaxation of selective pressure for some biological traits less impacted by cold adaptation. Specifically, the persistence of cold-stable water temperatures and high dissolved oxygen levels could be expected to relax selective pressures across biological systems that previously dealt with temperature and oxygen variability (Somero, 2010).

In addition to the influence of temperature, the isolation of the Southern Ocean has allowed the evolution of lineages characterized by substantial, and in some cases unique, physiological reorganization. This is exemplified by the members of the notothenioid family Channichthyidae (the icefishes) that are extraordinary for the absence of the respiratory pigment hemoglobin (Sidell and O’Brien, 2006). Oxygen is found solely in physical solution in icefish blood, and these fishes are thought capable of surviving in the Southern Ocean because the cold stable and well mixed Antarctic waters create a marine environment unusually rich with dissolved oxygen. While icefishes have undergone widespread and profound anatomical and physiological reorganization following the loss of hemoglobin, whether this is mirrored in large changes to the protein pool and corresponding genes or is mostly compensated through plasticity and gene expression regulation (Bargelloni et al, 2019) is less well understood.

Several prior studies have aimed to develop a broad understanding of the changes to protein-coding genes in the cryonotothenioids with polar evolution. Looking first at global biases in amino-acid composition that may explain adaptation to polar conditions, Berthelot et al. (2019) found evidence for only limited change. What they did find was an increase in leucine substitutions for methionine in the proteins of cryonotothenioids and hypothesized that these extra methionines had a role in redox regulation. This may be a particularly important defense for the Antarctic notothenioids as the oxygen-rich waters of the Southern Ocean would be expected to produce increased levels of reactive oxygen species that can be the source of macromolecule damage (Abele and Puntarulo, 2004).

Studies by Daane and colleagues (2020; 2019) have since investigated the genetic mechanisms that enable or constrain evolutionary change in the cryonotothenioids during their radiation in the Southern Ocean. Using the evolution of buoyancy adaptation among several Antarctic lineages (Eastman, 2020) as a model to explore the origins of novel traits in the cryonotothenioids, this was found to result from bone demineralization associated with developmental alterations similar to human skeletal dysplasia (Beck et al, 2021). However, the onset of positive diversifying selective pressure that reshaped these pathways were found to precede the origin of the cryonotothenioids, showing the role of historical contingency in shaping the capacity for adaptation in this group (Daane et al, 2019). Next, using the loss of oxygen transport genes as a model to explore constraints on genetic change from relaxation of purifying selective pressure in the cold Southern Ocean, Daane et al (2020) found that the pleiotropy of regulatory regions apparently served as a constraint on losses within transcriptional pathways. Additionally, this latter study found that the onset of relaxation of purifying selective pressure on pathways linked to oxygen transport was closely associated with the onset of freezing conditions in the Southern Ocean, contrasting with the earlier onset of positive diversifying selective pressure that enabled buoyancy adaptation among several cryonotothenioid lineages.

While results by Berthelot et al. (2019) as well as Daane and colleagues (2020; 2019) provide important insight into the timing and mechanisms of adaptive change in the cryonotothenioids, the suite of biological systems reshaped by evolution in the cold remains poorly surveyed in the cryonotothenioids. The aim of this study is therefore to survey the genes changed with evolution in chronic cold, and following the loss of hemoproteins, in order to gain a broad view of the biological systems that have been reshaped during the cryonotothenioid radiation. Towards this aim, we quantified the relative influences of positive and relaxed selective pressures during these physiological transitions, and identified the biological systems that have been altered through these changes.

## MATERIALS & METHODS

### Genomic resources, sequence, assembly, & filtering

To investigate how the pool of protein-coding genes have been changed by the ecological and physiological transitions experienced during the radiation of the cryonotothenioids, public genomic and transcriptomic resources were compiled for the cryonotothenioids along with a relevant background set of temperate and tropical fishes. For the cryonotothenioids, genomic-derived predicted peptides and their corresponding coding domain sequences (CDS) were only available at the time of this study for the three red-blooded species: *Dissostichus mawsoni* (Nototheniidae; Chen et al, 2019), *Notothenia coriiceps* (Nototheniidae; Shin et al, 2014), and *Parachaenichthys charcoti* (Bathydraconidae; Ahn et al, 2017). To extend this limited dataset, sequenced transcriptomic reads were downloaded for any species where more than one tissue had been sequenced, including the three red-blooded species with available genomic resources (Bargelloni et al, 2019; Berthelot et al, 2019; Bilyk et al, 2018; Kim et al, 2019; Song et al, 2019).

To provide a baseline for evaluating changes in selective pressure in the cryonotothenioids, predicted peptides and CDS were compiled for a background set of temperate and tropical fishes. These included two notothenioid temperate neighbors and sister species of cryonotothenioids, the Patagonian blenny *Eleginops maclovinus* (Eleginopidae; Chen et al, 2019) and *Cottoperca gobio* (Bovichtidae; Bista et al, 2020). The remaining species were selected from the teleost fishes available from Ensembl (Yates et al, 2020), with the aim of minimizing phylogenetic distance, avoiding obligatory freshwater species, avoiding species that inhabit freezing environments, and minimizing nested taxa. In total, protein-coding sequences were compared across 19 species in this analysis including 7 red-blooded cryonotothenioids, 5 hemoglobin-lacking icefishes, and 7 temperate and tropical fishes. The full list of the species used in this study and accession numbers for the genomic and transcriptomic resources used in the analysis can be found in the supplementary material (S Material 1).

The transcriptomic reads were then assembled for all of the cryonotothenioids. The reads were first cleaned with Fastp v0.20.1 (Chen et al, 2018) then assembled with Trinity v. 2.6.5 (Grabherr et al, 2011) using default parameters. Assemblies then went through preliminary filtering to remove redundancy with CD-hit v. 4.8.1 (Fu et al, 2012), and by removing contigs with less than 1 Transcripts Per Million (TPM) read coverage based on the transcriptome’s original sequenced reads. Predicted peptides and their corresponding CDS were then determined for the transcriptomes using TransDecoder v. 5.5.0 (Haas, 2021). These predicted peptides were subject to a final round of filtering meant to reduce much of the redundancy present in each species’ transcriptome, peptides were mapped against the Swissprot database using BlastP v2.10.1 retaining only the best match for each Swissprot accession number as determined by e-value. Finally, the relative completeness of species’ genome or transcriptome derived set of predicted peptides was evaluated using BUSCO v. 4 (Simão et al, 2015). In addition to providing a general measure of quality for each species’ gene sets, BUSCO scores were used to choose between genomic and transcriptomic derived predicted peptides and CDS when both were available for a given species.

### Phylogenetic reconstruction

To provide a phylogenetic frame of reference for later evolutionary hypothesis testing, we reconstructed the relationship among the 19 investigated species. This phylogeny was constructed using 2 mitochondrial (*16S, ND2*) and 11 nuclear genes (*MYH6, PKD1, SH3PX3, HECW2, SSRP1, PPM1D, RPS71, TBR1, PTR, RHO*, and *ZIC1*). Accession numbers for the genes used in this analysis are provided in S Material 2.

The sequences of each gene were aligned using MUSCLE v. 3.8.31 (Edgar, 2004), and the best-fit nucleotide substitution model of each gene was determined using the Akaike information criterion (AIC) in ModelTest-NG v0.1.6 (Darriba et al, 2020). The aligned sequences were concatenated and partitioned according to their best-fit models (GTR + I + gamma, mitochondrial: ND2 and 16S; GTR + I + gamma: MYH6 and PKD1; GTR + gamma: SH3PX3, HECW2, and SSRP1; GTR: PPM1D; HKY + gamma: RPS71, TBR1, PTR, and RHO; HKY + I: ZIC1). These substitution models were then implemented in Bayesian phylogenetic analyses using MrBayes v3.2.6 (Ronquist et al, 2012). The Markov chain Monte Carlo simulation was run for 100 million generations with four chains and sampled every 100 generations. MCMC convergence was assessed using the standard deviation of clade frequencies and potential scale reduction factor, and the first 25% of sampled trees were discarded as burn-in. Clade support was evaluated using posterior probabilities for nodes retained in the 50% majority rule consensus tree.

### Orthogroup inference, paralog pruning and building the multiple sequence alignments

Orthogroups were inferred across species from their filtered set of predicted peptides using OrthoFinder v. 2.5.1 (Emms and Kelly, 2019). The resulting orthogroups were then filtered to remove potential paralog contamination using a two-step process. First, PhyloTreePruner v. 1.0 (Kocot et al, 2013) was used to isolate the largest monophyletic subtree from the FastTree 2 (Price et al, 2010) generated gene trees produced for the contigs within each orthogroup. Second, the subtree was collapsed to a set of species-specific putative orthologs using BlastP against the *D. mawsoni* predicted peptide set. In this last step, contigs were only considered to have an orthologous relationship if they were best Blast hits, as determined by e-value, to the same *D. mawsoni* predicted peptide. If more than one contig from a species matched the same *D. mawsoni* predicted peptide then only the best blast hit, as determined by e-value, was retained. Only filtered orthogroups with representatives of all 19 target species were retained for constructing the multiple sequence alignments (MSAs).

Multiple sequence alignments were then generated for each retained orthogroup. The CDS for each orthogroup’s predicted peptides were codon aligned with GUIDANCE2 (Sela et al, 2015) using Prank v.140603 (Löytynoja, 2014), running 25 pseudo replicates per alignment. The alignments were then trimmed to remove any missing sites, and only alignments with all species and a minimum final length of 300nt were retained for evolutionary hypothesis testing.

### Identifying orthogroups under changed selective pressure

Orthogroups experiencing positive selective pressure were identified using the phylogeny-based strategy implemented in the Branch-Site Unrestricted Statistical Test for Episodic Diversification (BUSTED; Murrell et al, 2015) and the adaptive Branch-Site Random Effects Likelihood (aBSREL; Smith et al, 2015) methods in HyPhy (Kosakovsky Pond et al, 2020). The contrasting set of orthogroups experiencing a relaxation of selective pressure were identified using the (RELAX) method in HyPhy (Wertheim et al, 2015). All methods were implemented by specifying a set of foreground (or test) branches on which to test for positive or relaxed selection while the remaining branches were designated as background (or reference).

The orthogroups were first used to compare protein-coding genes in the red-blooded cryonotothenioids as foreground against the background set of temperate and tropical fishes to identify the set of orthologous genes that have come under changed selective pressure during the radiation of the cryonotothenioids in the progressively cooling Southern Ocean. Similarly, the icefishes were tested as foreground against a background of red-blooded cryonotothenioids to identify the set of orthogroups that have come under changed selective pressure specific to the loss of hemoproteins. A False Discovery Rate (FDR) adjusted p-value threshold of 0.1 was used in all tests to identify those orthogroups showing a significant change in selective pressure from the background set.

### Testing for functional enrichment

Gene ontology (GO) enrichment analysis was used to place the impacts of positive diversifying and relaxed selective pressures into biological context. The *D. mawsoni* predicted peptide set was annotated for GO terms using the Orthologous Matrix fast mapping utility (OMA browser; Altenhoff et al, 2021). GO terms were then applied to an orthogroup if it contained the original annotated *D. mawsoni* predicted peptide.

TopGO v. 2.44 (Alexa and Rahnenfuhrer, 2018) was used to test for functional enrichment among the isolated sets of genes under positive diversifying or relaxed selective pressures within the Biological Process (BP), Molecular Function (MF), and Cellular Component (CC) ontologies. Enrichment was tested using the sets of significant orthogroups identified by BUSTED, aBSREL, and RELAX, against a background set of 3452 orthogroups with representatives from all 19 species originally used in the analysis.

TopGO allows identification of enriched terms through over-representation enrichment using Fisher’s exact test as well as elimination and weighted methods that incorporate the GO hierarchy in order to limit redundancy in the final set of enriched terms. All three approaches were evaluated using a p-value threshold of 0.05 and enriched GO terms were assessed based on the literature of adaptive change to the Antarctic. From this comparison, the elimination algorithm was found to provide the most biologically informative set of enriched GO terms. The identified set of significant Biological Process GO terms was then clustered using the EnrichmentMap (Merico et al, 2010) plug-in of Cytoscape network visualization software v. 3.8.2 (Shannon et al, 2003).

## RESULTS

Metrics on the filtered genomic and transcriptomic resources used to identify orthogroups are presented in Fig 1, alongside the phylogenetic framework for evolutionary hypothesis testing (detailed in S Material 3). Additional information on the tissue content of each transcriptome is presented in S Material 4 and comparisons between the genomic and transcriptomic derived predicted peptides for species, where both were available, are presented in S Material 5. In total, 3,453 orthogroups were identified that contained all species in the 19-taxa dataset. All of these orthogroups were used for alignment construction then evaluated using HyPhy to identify genes showing signatures of either positive or relaxed selective pressures.

**Figure 1.**
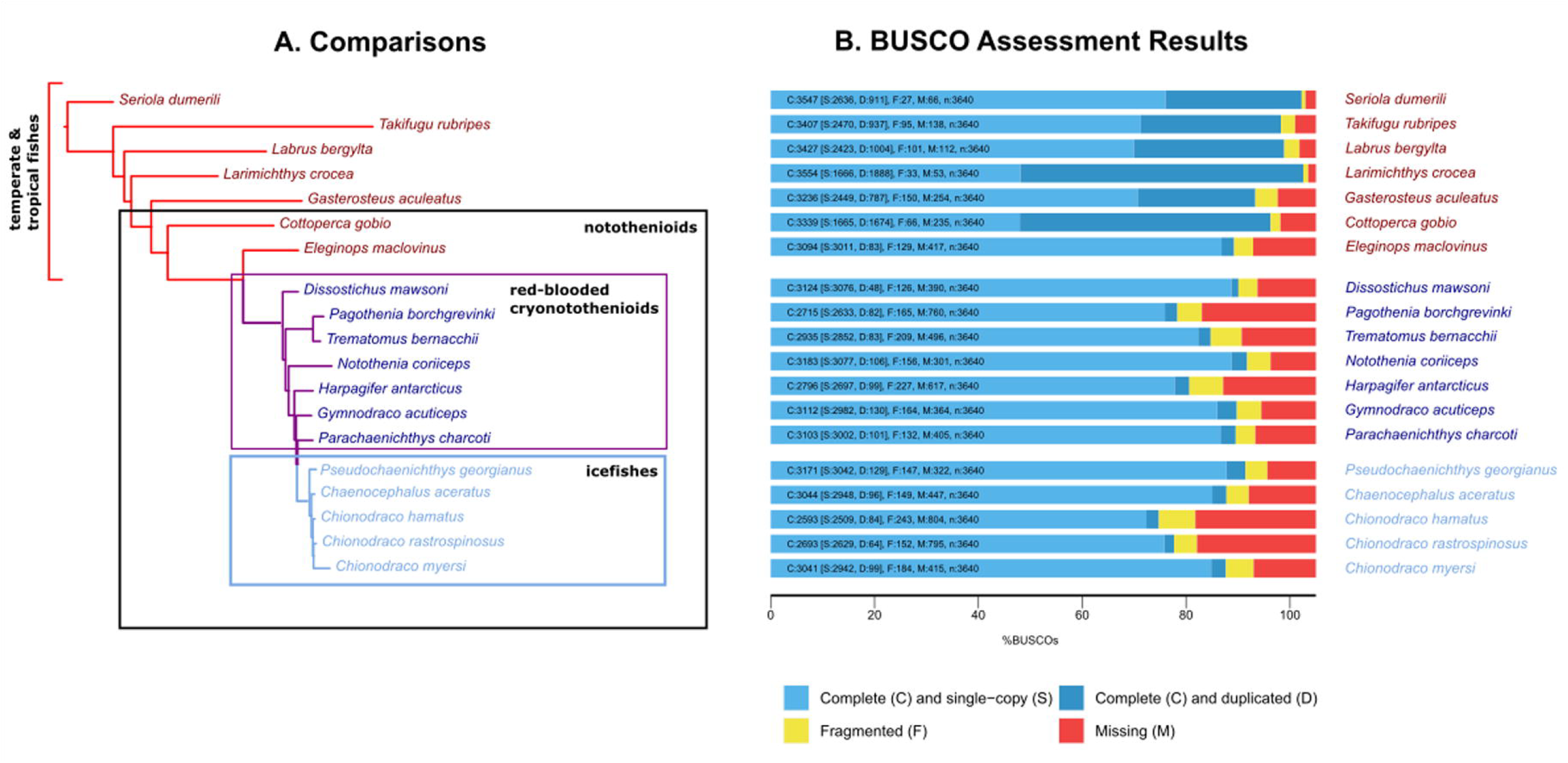
Panel A shows the comparative groupings used in the analysis, with species in red highlighting temperate/tropical fishes, species in purple highlighting the red-blooded cryonotothenioids, and species in blue highlighting the icefishes. Panel B highlights BUSCO metrics for the peptide sets used for original ortholog detection in all 19 target species.

First, the red-blooded cryonotothenioids were compared to temperate and tropical fishes to identify changes related to evolution in the chronic cold. From this comparison, aBSREL and BUSTED identified 113 and 89 orthogroups respectively under positive diversifying selection, with 160 distinct orthogroups identified as experiencing positive selective pressure from either of the two analyses (Fig 2A). RELAX then identified 183 orthogroups under relaxed selective pressure (Fig 2A).

**Figure 2.**
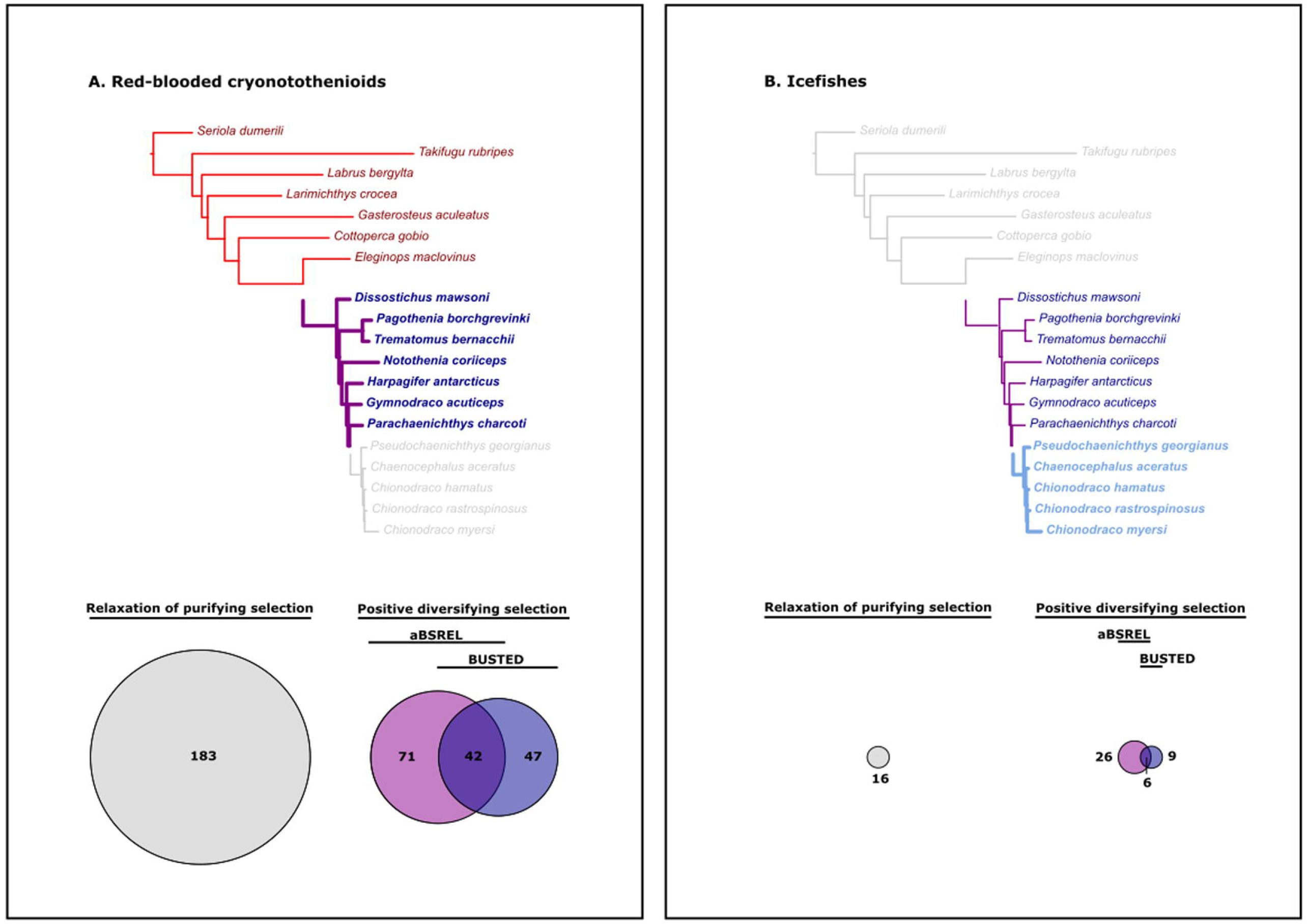
Number and distribution of orthogroups showing changes in selective pressure with the transition to freezing water temperatures and with hemoprotein loss. The phylogenetic trees show the comparison each analysis is based on while the Venn diagrams show the number of genes identified by RELAX as being under relaxation of purifying selective pressure, or by aBSREL and BUSTED as being under positive diversifying selective pressure.

Next, the icefishes were compared against a background set of the red-blooded cryonotothenioids, to identify genes under changed selective pressure during the diversification of icefishes and with hemoprotein loss. This comparison provided a far weaker signal of changed selective pressure, for both positive and relaxed selective pressures. The signature of significant positive selective pressure was identified on only 15 orthogroups using BUSTED, and in 32 orthogroups with aBSREL, resulting in a total of 41 distinct orthogroups identified as experiencing positive selective pressure from either analysis (Fig 2B). Similarly, relaxed selective pressure was detected for only 16 orthogroups using RELAX (Fig 2B).

To put the sets of orthogroups showing signatures of changed selective pressure into biological context, we tested for functional enrichment of genes showing signatures of changed selective pressure by GO enrichment analysis. Looking at the red-blooded cryonotothenioids, there were 36 total enriched GO terms from the set of genes identified under positive diversifying selective pressure by either aBSREL or BUSTED (S Material 6-9). These included 30 terms in the BP ontology, 4 terms in the MF ontology, and 2 terms in the CC ontology (S Material 9). Focusing on the Biological Process ontology, clustering identified seven clusters of GO terms connected by shared sets of genes (Fig 3A). These had broad functional impacts on regulation of biosynthetic processes, cell cycle progression, protein polymerization, cell adhesion, endopeptidase activity, the response to temperature, blood circulation, as well as smaller sets related to the response to organic cyclic compounds and lipoprotein metabolic processes.

**Figure 3.**
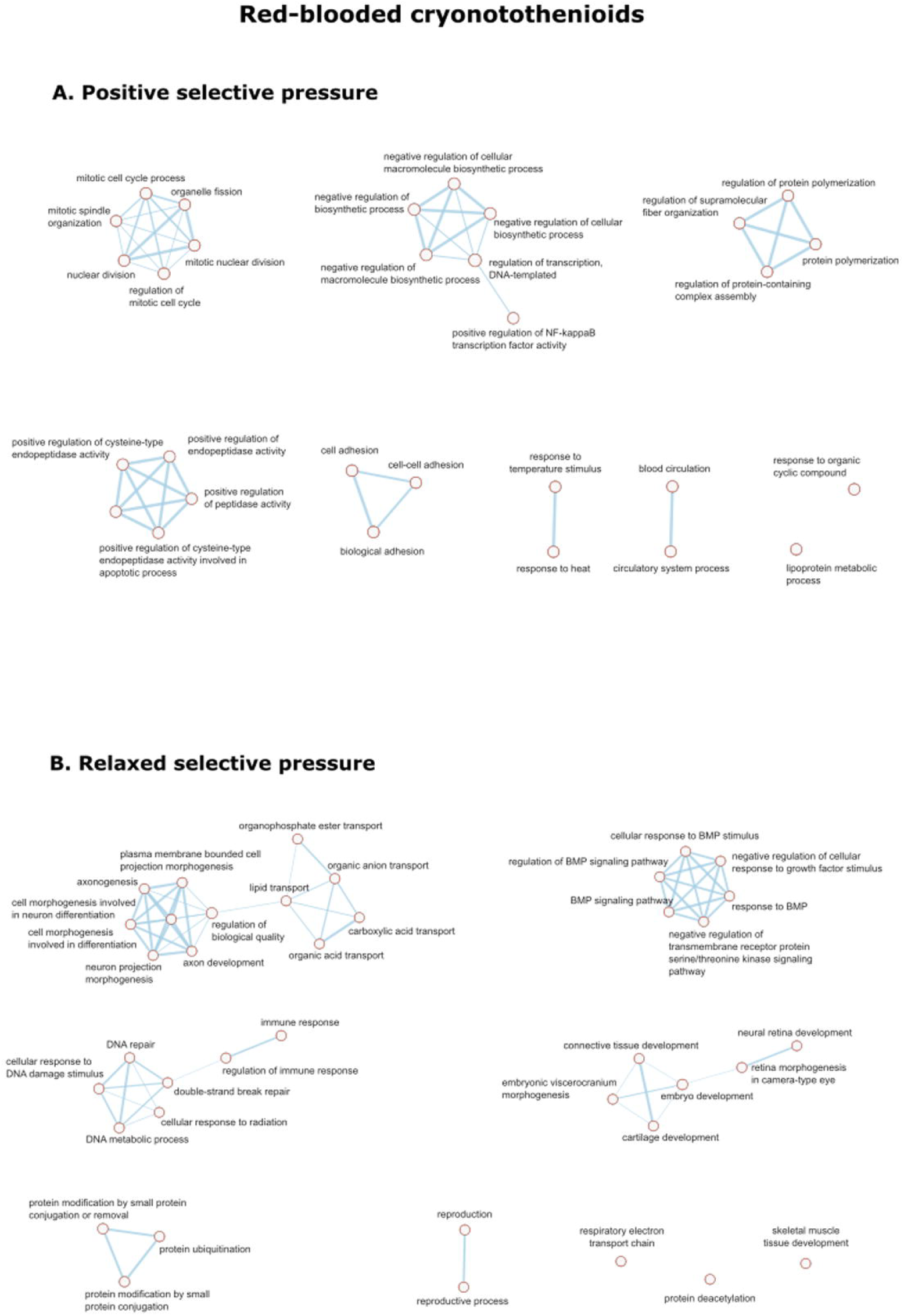
GO terms enriched in the Biological Process ontology among the red-blooded cryonotothenioids among genes showing signatures of positive diversifying and relaxed pervasive selective pressures.

Looking at orthogroups showing signatures of relaxed purifying selective pressure in the red-blooded cryonotothenioids, these included 39 terms in the BP ontology, 15 terms in the MF ontology, and 2 terms in the CC ontology (S Material 10). Within the Biological Process ontology, clustering identified six clusters of GO terms connected by shared sets of genes (Fig 3B). These show functional impacts on the regulation of biological quality, Bone Morphogenetic Protein (BMP) signaling, connective tissue development, DNA repair, protein ubiquitination, reproduction, the respiratory electron chain, protein deacetylation, and skeletal muscle tissue development.

Despite the small number of genes identified with significant changes in selective pressure in the icefishes relative to the red-blooded cryonotothenioids, we did continue to identify enriched GO terms in the icefish. There were 31 enriched GO terms from the set of genes identified under positive diversifying selective pressure by either aBSREL or BUSTED (S Material 11-13). These included 24 terms in the BP ontology, 4 terms in the MF ontology, and 3 terms in the CC ontology (S Material 13). Clustering identified two modules of GO terms from the BP ontology connected by shared sets of genes (Fig 4A), with most positively selected orthogroups in a large, multifunctional module. The remaining positively selected genes belonged to a smaller module centered on protein modification. Looking at orthogroups showing signatures of relaxed selective pressure, these included 13 terms in the BP ontology, 11 terms in the MF ontology, and 2 terms in the CC ontology (S Material 14). Within the Biological Process ontology, clustering identified two modules of GO terms from the BP ontology connected by shared sets of genes (Fig 4B), with most belonging to a module related to inorganic ion transport and the remainder focused on cell cycle control.

**Figure 4.**
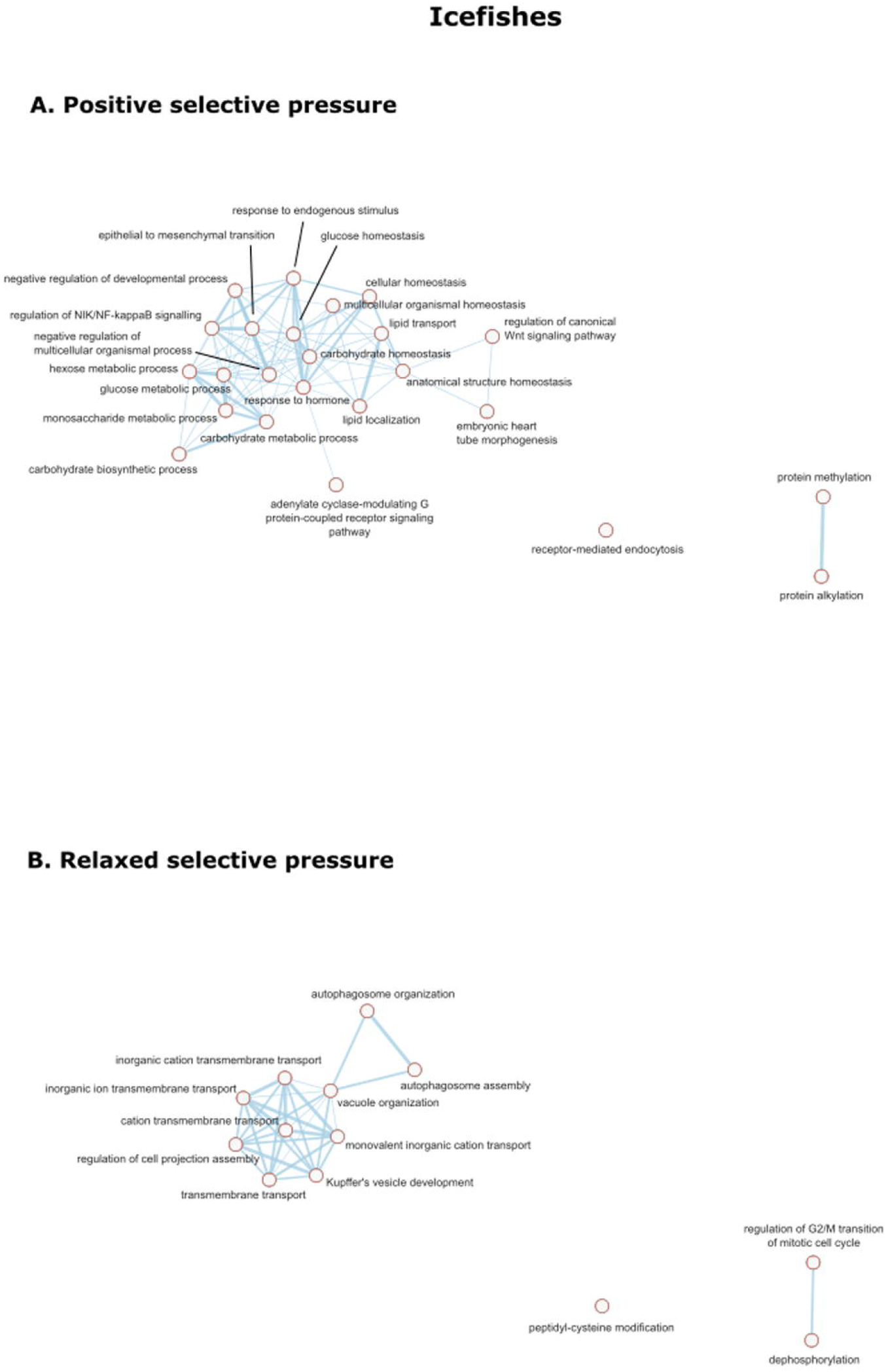
GO terms enriched in the Biological Process ontology among the icefishes among genes showing signatures of positive diversifying and relaxed pervasive selective pressures.

## DISCUSSION

While the radiation of the cryonotothenioids in the isolation of the Southern Ocean has exposed them to substantially different selective pressures compared to temperate and tropical fishes, it has remained unclear how this has acted across the cryonotothenioid’s biological systems. Here, we find that evolution in the chronic cold is correlated with widespread signatures of both positive and relaxed selective pressures. This is in contrast with the loss of hemoproteins in the icefishes, where few genes showed signatures of changed selective pressure compared to the red-blooded cryonotothenioids.

### Genes under positive diversifying selective pressure in the red-blooded cryonotothenioids

The changes among protein-coding genes in the red-blooded cryonotothenioids provide a window into the challenges to maintaining the functions of life in the Southern Ocean’s freezing and oxygen-rich waters. By comparison against temperate and tropical species, genes under positive selection in the red-blooded cryonotothenioids serve to identify those particular biological functions with essential roles in polar adaptation. Here, we found that positive diversifying selective pressure acted on genes related to biosynthetic processes, cell cycle progression, protein polymerization, cell adhesion, and endopeptidase activity, with smaller sets of genes related to temperature response and blood circulation (Fig 2A).

Prominent among these were the set of genes with roles in **biosynthetic processes**. Prior studies have suggested that folding of nascent macromolecules and their subsequent conformational maintenance are challenges to life in freezing polar waters (Place et al, 2004), but here we have found a previously unappreciated signal of positive selection on the machinery of transcription. These included genes that mediate RNA polymerase activity (*MED23* and *TCEA1*) and mRNA processing (*THOC5*). Transcription shows reduced efficiency and increased errors as an organism moves from its preferred temperature towards temperature extremes (Meyerovich et al, 2010). This would suggest that genes related to transcription would come under selection to ensure continued accurate gene expression with temperature adaptation to new thermal environments, such as the transition from temperate to polar conditions. Several genes related to the process of translation were also found under positive diversifying selective pressure. Notably though, these were specific to translation in the mitochondria including genes involved in mitochondrial ribosomal biogenesis (*UTP15*) as well as components of the mitochondrial ribosomes themselves (*MTG2* and *MRPL13*).

The set of **biosynthetic process** genes under selection also included a suite of transcription factors, transcription repressors. These include genes that regulate cell growth and proliferation (*BAP1, KAT8, PDCD4, ZGPAT*), metabolism (*USF1*), RNA polymerase III activity (*MAF1*), and the cellular response to stress (*CREB1*), suggesting these are all key challenges to proper cellular function at freezing temperatures. A set of genes associated with immune and inflammation responses were also among this set under positive diversifying selection including genes with broad roles in the immune system (*STAT1*), with roles regulating innate (*ELF4, IRF3*) and adaptive immunity (*NFATC3, RFX1*), as well as several genes within the NF-KB pathway (*IKBKB, NFKBIA*). However, the role that selection from a polar environment has had in reshaping immune function remains unclear as a number of immune regulatory genes were prominently among those under relaxed purifying selective pressure.

Signatures of positive selective pressure were seen among genes related to **cell cycle progression**, including genes with a direct role in its regulation and progression (*CC14A, GNAI1, PDC6I, PRCC*). Cold Antarctic water temperatures of the Southern Ocean act to generally slow cellular processes, including progression of the cell cycle creating challenges to growth and development of local organisms. While little studied, prior work comparing cell cycle progression between the cryonotothenioid *Harpagifer antarcticus* and its sub-Antarctic congener *H. bispinis* has suggested cold compensation of the cell cycle in cryonotothenioids as an adaptation to limit this slowing influence of low temperatures (Brodeur et al, 2003). Selection on cell-cycle genes further supports the assertion of adaptation in the control and mechanisms of that underlie progression and functioning of the cell cycle. On functional adaptation, signatures of positive selection were observed on genes that act to maintain the stability of microtubules during mitosis (*NUF2, DRG1*) and with a role in chromosome movement (*KIF22*). Beyond their role in supporting the actions of the cell cycle, selection on this latter set of genes appears to reflect a general trend of selection on protein polymerization, especially cytoskeletal elements.

Decreases in temperature can impede **protein polymerization** and stability of cytoskeletal elements (Li and Moore, 2020), and adaptive change to enable efficient polymerization at low temperature has long been recognized among these proteins in the cryonotothenioids (Detrich et al, 2000). Here, we find that such signatures of adaptation appear to extend beyond the elements themselves and into a wider number of genes that support their function including two genes with roles promoting polymerization among cytoskeleton elements (*DRG1, PROF2*). Similarly, genes related to **cellular adhesion** were also found to be under positive diversifying selective pressure. These included components of the junctions themselves (*CADH1, DSC3*), that link intercellular connections to the cytoskeleton (*CTNA1*), and that regulate cell junction organization (*TMM47*). Not previously noted as an adaptive change among the cryonotothenioids, cell-cell linkages may generally be challenged to organize and maintain adhesion in the cold (Zieger et al, 2011), and thus have come under selective pressure with evolution in the cold in order to maintain the integrity of tissues.

Several genes that **regulate endopeptidase activity** (*CFLAR, LCK, PDCD6*) were under positive selection. Though the observed genes are varied in their biological roles, they were all linked as regulators of apoptosis pathways. Change in the behavior of apoptosis pathways with cold adaptation has been suggested by comparative transcriptomic studies looking at model fish species (Hu et al, 2016). This may suggest change to the functioning of genes cell death pathways is also associated with the evolution of cold hardiness.

Among smaller sets of genes, positive diversifying selective pressure had functional influences on the **response to heat** and on **blood circulation**. While the cold-stable water temperatures of the Southern Ocean would be expected to primarily relax selective pressure on heat responding genes, positive selective pressure was seen acting on several including a temperature sensor (*TRPV1*), in addition to genes with supporting roles within the heat shock response (*PDCD6*), and on a gene with a direct role in responding to environmental stress (*GSH1*). Of the three, the evolution of temperature sensors is little studied in polar animals but these likely represent a class of genes labile to rapid evolutionary change with temperature adaptation (Somero, 2010). Genes with a role in blood circulation showed signatures of selection, including genes with a role in blood pressure (*EDNRA, MYZAP*) that may be among the suite of adaptations that allow the cryonotothenioids to overcome increased blood viscosity at low temperature (Egginton and Rankin, 1998).

Finally, signatures of positive diversifying selective pressure were seen in genes that **respond to organic cyclic compounds** and **lipoprotein metabolic processes**. Genes responding to organic cyclic compounds include subunits of guanine nucleotide-binding proteins (*GNAI1, GNB5B*) which may signify cold-adaptation to maintain the basic function of essential intracellular signaling mechanisms. A variety of genes with roles in lipoprotein metabolic processes are under selection in the red-blooded cryonotothenioids (*AB17A, APOA1, ZD18A, ZD23B*). Compared to temperate fishes, the cryonotothenioids have larger proportions of polyunsaturated lipids in their membranes and rely more heavily on lipids as an energy source (Todgham et al, 2017), both suggesting that lipid biosynthetic pathways have come under selective pressure.

### Genes under relaxed selective pressure from chronic cold-water temperatures in the red-blooded cryonotothenioids

The chronic nature of cold-water temperatures in the Southern Ocean would itself have been an important influence on protein evolution in the cryonotothenioids just like the freezing temperatures of these waters. While the cold would impose selective pressure for cold-hardiness on biological systems, the stability of these low temperatures would be expected to relax purifying selective pressure, especially among biological systems that respond to thermal variability. However, we found the signature of relaxed purifying selective pressure across a wider set of biological functions including the regulation of biological quality, BMP signaling, connective tissue development, DNA repair, protein ubiquitination, and reproduction. Additionally, smaller sets of genes were associated with the respiratory electron transport chain, protein deacetylation, and skeletal muscle development. Ultimately these seem to suggest two central roles of relaxed selective pressure, first across a suite of biological systems that help maintain homeostasis, and second across a suite of genes that pattern morphology.

Prominent among genes under relaxed selective pressure were those associated with the **regulation of biological quality**. This gene set appears to mirror the division seen among the broader set of genes showing signatures of drift, between genes with roles in homeostatic processes and others with roles shaping morphology. For the former, signatures of relaxed selective pressure were found among genes in a number of biological systems that maintain homeostasis and that may now be under reduced demand given the stability water temperatures and high oxygen concentration of Southern Ocean waters. These include iron homeostasis (*FBXL5, SLC40A1*), mitochondrial function (*FAKD1, FAKD3, PDK3, PRELI*), platelet activation (*PAR1, VWF*), DNA damage response (*NIPLB*), oxidoreductase activity (*RDHE2, RD10A*), integrin-linked kinase signaling (*PARVB*), apoptosis (*CAS9*), and an energy carrier (*GTR1*). Besides these homeostasis-related genes, a second theme seen in the genes under relaxed selective pressure were changes that appear linked to the morphological diversity that has developed in the cryonotothenioids. These include genes with roles associated with axonogenesis (*CASP9, DREB, CYFP2, KIF5B*) which, when combined with the signature of positive selection on microcephalin (*MCPH1*), may show the influence of pedomorphism on cranial development in the cryonotothenioids (Albertson et al, 2010), as cranial development and brain size are linked (Koyabu et al, 2014).

The diverse range of systems affected by relaxation selective pressure in the Southern Ocean is seen among a group of genes that carry out **protein modification by small protein conjugation**, a common mechanism of regulating cellular activity. These include genes that regulate the cell cycle (*CDC34, OBI1*), the DNA damage response (*ARI1, DDB2, UBP7*), iron homeostasis (*FBXL5*), tRNA biosynthesis (*CTU1*) and neurogenesis (*DTX1, UBA6*). Additionally, several genes with direct roles in the recycling of misfolded proteins showed relaxation of purifying selective pressure, including a ubiquitin activator (*UBA6*), as well as a ubiquitin-conjugating enzyme directly involved in endoplasmic reticulum-associated degradation (*UBE2K*). The latter may result from the loss of inducibility in the endoplasmic reticulum stress response (Bilyk et al, 2018), which would presumably reduce peak demand on endoplasmic reticulum-associated degradation system.

Signatures of relaxed selective pressure were seen among genes engaged in **DNA repair**. These span the acute response to DNA damage (*PARP3, XRCC6, DDB2, R51A1*), repair that occurs during replication (*MSH2, SAMH1*), normal DNA replication (*KIF22, DNMT1*), as well as DNA damage driven apoptosis (*CASP9*). The relaxation of selective pressure on this set of genes maintaining DNA integrity may be the result of the same reduction in thermal variability that allowed for the loss of the heat shock response in the cryonotothenioids (Hofmann et al, 2000). Heat shock and thermal variability are key sources of DNA damage, both as sources of acute damage, and as a cause of errors during the replication of DNA (Kantidze et al, 2016). The loss of this thermal variation could thus remove an important source of DNA damage and therefore lead to potential atrophy of damage responding pathways. Besides thermal variation, this relaxation of selective pressure may alternatively result from the slower metabolism and cell cycle observed in the cryonotothenioids both of which would be expected to naturally limit the incidence of DNA damage. While high production of ROS in oxygen-rich waters is thought to increase the risk of DNA damage in the Southern Ocean, this may be a lesser risk to DNA integrity compared to thermal variability, or it may be mitigated through modulation of gene expression (Abele and Puntarulo, 2004; Chen et al, 2008).

While genes with roles in the development and functioning of the immune system were found among the set of biosynthetic process genes under positive selective pressure, another set of genes with roles in the **immune response** were found under relaxed selective pressure. These included genes with roles in the differentiation of immune cells (*NR4A3, KPCL*), components of the membrane attack complex (*CO8A*), host restriction factors (*SAMH1*), mediating NF-kappa-B signaling (*ERBIN*), and interleukin signaling (*STAT6*). Ultimately the competing signatures of selection in the immune system appear to match inconclusive measures of change in empirical investigations of immune function in Antarctic fishes. Prior work has shown continued transcriptional response of immune genes following immune stimulation (Ahn et al, 2016; Buonocore et al, 2016), though when challenged by lipopolysaccharide injection, the cryonotothenioids *N. coriiceps* and *N. rossii* showed a less than robust immune response marked by considerable individual variability (Ahn et al, 2016). While the immune system in cryonotothenioids appears to be under selective pressure, it remains unclear exactly how the selective forces of the Southern Ocean may have altered its behavior.

Relaxation of selective pressure was seen among genes of the mitochondria’s **respiratory electron transport chain** that is responsible for transferring electrons from an electron donor to electron acceptors driving the process of oxidative phosphorylation. This change in selective pressure was detected in several components of the electron transport chain itself (*SDHA, QCR1, NDUA8, COX41*), amid signatures of relaxed selective pressure on other genes related to mitochondrial function detected under other GO terms (*COA7, NDUA8, PRLD1*). Similar results were obtained in a previous study for mitochondrially-coded genes for which a prevalence of relaxation of the selective pressure in comparison with the non-Antarctic notothenioids was detected (in particular for *COX3, COB, NAD2, NAD4*, and *NAD6*; Papetti et al, 2021). While this may be a result of increased oxygen saturation in cold Antarctic waters, this trend towards relaxation of selective pressure may also be a result of the reduction in resilience to thermal variability in these systems as suggested in prior work (Mark et al, 2012) and may have persistently influenced the inception of the evolution of cryonotothenioids.

Genes falling under the heading of **reproduction and reproductive process** do include some related to early stages of development (*SPIN1, CXCR4*) as well as a small number of genes with broader roles in cell cycle progression (*GCP3, KINH, NUF2*). Early development in polar marine invertebrates is far slower than related temperate or tropical species (Peck, 2016), and long-lasting larval stage is seen in the cryonotothenioids as well (Kellermann, 1989). The relaxation of selective pressure in development genes may reflect this extended development process. One can also speculate that this extension of early developmental stages and paedomorphism may both be part of a suite of ecological adaptations that have enabled cryonotothenioids to colonize large areas of the Antarctic shelf and several ecological niches that opened in the freezing Southern Ocean.

The genetic signatures of morphological diversity are also seen in a set of genes associated with BMP signaling and connective tissue development. **BMP signaling** plays important roles in embryogenesis and development in addition to its role in bone formation (Wang et al, 2014). Here, we see the influence of relaxed selective pressure primarily on genes involved in central nervous system development (*SPART, ACATN*). Alongside the relaxation of selective pressure previously noted among axonogenesis-related genes, the changes in BMP signaling and connective tissue development genes are likely part of the suite of genetic changes that enable the diversity of skull morphology now seen among the cryonotothenioids.

Relaxed selective pressure on genes with roles in **connective tissue development** similarly appears to be part of the genetic changes behind the ecological and morphological diversification of the clade. As with the changes in BMP signaling, signatures of change were seen in genes associated with craniofacial development (*DCAF7, CRLD2, SIAS*) which may relate to paedomorphism on cranial development (Albertson et al, 2010; Koyabu et al, 2014). However, the largest set of connective tissue genes were involved in chondrocyte differentiation and skeletal development (*SOX9, KINH, TEAD1*), or with roles in bone degeneration (*CSF3R*). These would likely be part of the suite of genetic changes behind the reduction in bone density in these fishes that has allowed the evolution of neutral buoyancy in some lineages (Eastman, 2020).

### Genes under changed selective pressure in the icefishes

Unlike the transition from temperate to polar environments, the loss of hemoproteins appears to have shifted selective pressure on a much smaller suite of genes. The limited scale of change among the tested orthogroups was consistent for both positive and relaxed selective pressures. While few genes showed significant signs of change, several biological themes could still be identified.

Positive selective pressure was found to act broadly, corresponding to several known areas of physiological and morphological change in the icefishes. These included genes with roles in cardiac morphogenesis, myogenesis, and vascular development (*MEF2C, NIPLB*) and correspond to the highly modified cardiovascular system seen in the icefishes that allows the larger blood volume necessary to transport oxygen in blood plasma without the presence of red-blood cells (Sidell and O’Brien, 2006). Interestingly, Bargelloni et al. (2019) found that several genes of the Myocyte enhancer factor (MEF family), including *MEF2C* are largely upregulated in the icefish and these may play an important role in the larger mitochondrial density seen in icefish cells. Similarly, changes in energy metabolism genes (*F16P1, GTR1, PAQR1*) correspond to increases in mitochondrial size and density within icefish cells, and perhaps the need to keep cellular metabolism fueled in these fishes as oxygen availability is more limited. Selection was seen in genes related to cartilage biosynthesis (*CHSTB*) and neural patterning (*IRX3*) both of which would likely be tied to the continued modification of skeletal and skull shape than what is seen in the remaining cryonotothenioids. These were in addition to further changes seen in developmental pathways (*PRMT8*), signal transduction (*GBB1, LPAR4*), and the regulating gene expression (*PAGR1*). Relaxation of selective pressure appears to have had an even smaller influence among tested genes. Changes were detected on a varied group of genes with roles in transmembrane transport (*AQ7, KCNK5, VATF*), autophagosome organization (*RBCC1*), mitotic cell cycle progression (*MPIP1*), along with more general roles in the cell (*PLPP3*).

While there continue to be signatures of genes under both positive and relaxed pressure in the icefishes compared to the red-blooded cryonotothenioids, it is notable that changes among the pool of protein genes are so much smaller than what is observed between temperate and polar species. Given that the icefishes show widespread anatomical and physiological modifications compared to the red-blooded cryonotothenioids, why might these show such muted signatures of genetic change?

One possibility is that the relatively recent divergence of the icefishes has not allowed sufficient time to see widespread change in protein coding genes even if selective pressures have shifted. Alternatively, Bargelloni et al. (2019) suggested that the loss of hemoglobins has been accompanied by compensation mechanisms like silencing or gene expression regulation more than positive or relaxed selection to maintain the additional function of genes originally involved in erythropoiesis. Finally, prior work has also shown that the oxygen-rich Southern Ocean environment relaxed selective pressure on oxygen carriers before hemoprotein loss is observed in the icefishes, as seen in the reduction in hemoglobin isoform complement (Eastman, 1993) and in the attenuation of regulatory elements upstream of hemoglobin (Lau et al, 2012) both in the red-blooded cryonotothenioids. As a result, the icefishes may simply reflect the same selective forces exerted across the cryonotothenioids as a whole. Recent work by Daane et al (2020) suggested as much, finding convergent signatures of drift in anemia-related genes in several red-blooded notothenioid lineages as well as the icefishes. This would be consistent with our view of reduced signatures of selective pressure in the protein-coding pool in the icefishes.

## CONCLUSIONS

Our study shows that the genomes of the cryonotothenioids have been shaped almost equally by positive diversifying and relaxed purifying selective pressures in the chronically cold and oxygen-rich Southern Ocean, making these fish strongly adapted to and ecologically dependent on the polar environment. While recent work has emphasized the roles of plasticity or pleiotropy as agents that may have limited the scope of selective change in the cryonotothenioids, evolution in cold-stable water temperatures was still found to have exerted change in genes across a wide range of biological functions. This study has further highlighted the pivotal role of genes related to mitochondrial function, biosynthesis, protein polymerization, and skeletal development in the evolution and diversification of the cryonotothenioids, some unique and new genetic changes have also emerged.

Positive selective pressure was most strongly felt on genes with roles in biosynthetic processes, with novel changes identified among the machinery of transcription, in control over cell cycle progression, and in cell adhesion. Contrasting with this, relaxation of selective pressure exerted an influence on several homeostatic processes, as the chronic nature of cold temperatures and high oxygen concentrations reduced the demand on biological systems that once dealt with their variability. However, relaxation of purifying selective pressure also appears to have strongly influenced many genes with roles in CNS and skeletal development, suggesting this has been a critical force in generating the morphological and ecological diversity now seen in the cryonotothenioids. Finally, in icefishes, the loss of hemoglobin has probably impacted less the genomic variability via selective pressure while possible alternative mechanisms have been put into action to compensate for the unique loss and to maintain function.

The Antarctic fish have been exposed to long and intense selective pressure for survival in freezing waters, and are characterized by long development and generation times, deferred maturity, and extended life spans (Peck 2018). Since all changes highlighted in this study are the result of long-term exposure to cold, and have led to profound changes in the cryonotothenioid genome, it seems very unlikely that any reverse adaptation might be possible in the short term of the predicted climatic alteration occurring in Antarctica (Masson-Delmotte et al, 2021).

## ACKNOWLEGMENTS

The authors would like to thank Dr. Agnès Dettaï (Muséum national d’Histoire naturelle, Paris, France) and Enrico Negrisolo (University of Padova, Padova, Italy) for advice on the phylogenetic markers used in this analysis. The authors would further like to thank Katherine R. Murphy for her comments on the manuscript. Many of the selection test methods applied in this study were facilitated using best practices and scripts provided as part of a Next Generation Sequencing-based workshop sponsored by the National Science Foundation (Award # 1744877 to Scott Santagata, Biology Department, Long Island University, New York, USA).

## AUTHOR CONTRIBUTIONS

KTB and CP developed the original comparison between temperate and polar species, KTB and XZ developed the comparison between the icefishes and red-blooded cryonotothenioids. XZ built the phylogenetic framework for the study and KTB carried out the selective pressure analyses. All three authors contributed to the analysis of the selective pressure results and writing the manuscript.

## DATA AVAILABILITY STATEMENT

All genomic and read data used in this study were downloaded from publicly available repositories as detailed in the supplementary material.

## SUPPLEMENTARY MATERIAL

S Material 1: Genomic and transcriptomic resources used in this study

S Material 2: Genes used in Phylogenetic Reconstruction

S Material 3: Phylogenetic tree used in selective pressure analyses

S Material 4: Tissue completeness of each notothenioid species’ transcriptome

S Material 5: Comparison of genomic and transcriptomic derived predicted peptides using BUSCO for three cryonotothenioid species

S Material 6: Enriched GO terms for orthogroups identified under positive diversifying selective pressure by either (union) BUSTED or aBSREL in the red-blooded cryonotothenioids

S Material 7: Enriched GO terms for orthogroups identified under positive diversifying selective pressure by only BUSTED in the red-blooded cryonotothenioids

S Material 8: Enriched GO terms for orthogroups identified under positive diversifying selective pressure by only aBSREL in the red-blooded cryonotothenioids

S Material 9: Enriched GO terms for orthogroups consistently identified under positive diversifying selective pressure by both (intersection) BUSTED and aBSREL in the red-blooded cryonotothenioids

S Material 10: Enriched GO terms for orthogroups consistently identified under relaxed selective pressure by RELAX in the red-blooded cryonotothenioids

S Material 11: Enriched GO terms for orthogroups identified under positive diversifying selective pressure by either (union) BUSTED or aBSREL in the icefishes

S Material 12: Enriched GO terms for orthogroups identified under positive diversifying selective pressure by only BUSTED in the icefishes

S Material 13: Enriched GO terms for orthogroups identified under positive diversifying selective pressure by only aBSREL in the icefishes

S Material 14: Enriched GO terms for orthogroups consistently identified under relaxed selective pressure by RELAX in the icefishes

