## Supplemental Material for "Positive and relaxed selective pressures have both strongly influenced the evolution of cryonotothenioid fishes during their radiation in the freezing Southern Ocean"

**Supplementary Material**

**Table of Contents**

**SECTION**	**PAGE**

S Material 1 …	1

S Material 2 …	4

S Material 3 …	6

S Material 4 …	7

S Material 5 …	8

S Material 6 …	9

S Material 7 …	10

S Material 8 …	12

S Material 9 …	13

S Material 10 …	14

S Material 11 …	16

S Material 12 …	17

S Material 13 …	18

S Material 14 …	20

**S Material 1. Genomic & Transcriptome sequences used in the data analysis**

| **SPECIES** | **ACCESSION** | **TISSUE** |
| --- | --- | --- |
| *Eleginops maclovinus* (genome) ^x^ | doi:10.5524/102163 | N/A |
| *Eleginops maclovinus* (transcriptome) | SRR5210465  (pool) | Brain  Eye Cup  Gill  Head Kidney  Heart  Intestine  Liver  Lens  Spleen  Stomach |
| *Dissostichus mawsoni* (genome) ^x^ | doi:10.5524/102162 | N/A |
| *Dissostichus mawsoni* (transcriptome) | SRR6794060 | Brain |
|  | SRR6794071 | Caudal Kidney |
|  | SRR6794062 | Gill |
|  | SRR6794059 | Head Kidney |
|  | SRR6794061 | Liver |
|  | SRR6794069 | Ovary |
|  | SRR6794065 | Pelvic Girdle Bone |
|  | SRR6794063 | Red Muscle |
|  | SRR6794066 | Skin |
|  | SRR6794070 | Spleen |
|  | SRR6794068 | Small Intestine |
|  | SRR6794067 | Stomach |
|  | SRR6794064 | White Muscle |
| *Notothenia coriiceps* (genome) ^\|^ | GCF_000735185.1 | N/A |
| *Notothenia coriiceps* (transcriptome) | SRR1015840 | Blood (control) |
|  | SRR1015879 | Blood (heat stress, 48 hours) |
|  | SRR1015880 | Blood (heat stress, 48 hours) |
|  | SRR1015881 | Blood (cold stress, 48 hours) |
|  | SRR1015882 | Blood (cold stress, 48 hours) |
|  | SRR1015883 | Brain (control) |
|  | SRR1015884 | Brain (control) |
|  | SRR1015885 | Brain (heat stress, 48 hours) |
|  | SRR1015886 | Brain (heat stress, 48 hours) |
|  | SRR1015887 | Brain (cold stress, 48 hours) |
|  | SRR1015888 | Brain (cold stress, 48 hours) |
|  | SRR1015889 | Blood (cold stress, 48 hours) |
|  | SRR1015890 | Blood (cold stress, 48 hours) |
|  | SRR1015891 | Skin (heat stress, 48 hours) |
|  | SRR1015892 | Skin (heat stress, 48 hours) |
|  | SRR1015893 | Skin (cold stress, 48 hours) |
|  | SRR1015894 | Skin (cold stress, 48 hours) |
|  | SRR1015895 | N/A |
|  | SRR1015896 | N/A |
|  | SRR1015897 | N/A |
|  | SRR1015898 | N/A |
|  | SRR1015899 | N/A |
|  | SRR1015900 | N/A |
|  | SRR1015901 | N/A |
| *Trematomus bernacchii* (transcriptome) | SRR7164563 | Gonad, Male |
|  | SRR7164560 | Head Kidney |
|  | SRR7164559 | Heart |
|  | SRR7164558 | Liver |
|  | SRR7164565 | Muscle |
|  | SRR7164561 | Skin |
|  | SRR7164570 | Spleen |
|  | SRR7164564 | Stomach |
| *Pagothenia borchgrevinki* (transcriptome) | SRR5210464  (pool) | Brain  Eye Cup  Gill  Head Kidney  Heart  Intestine  Lens  Liver  Spleen  Stomach |
| *Harpagifer antarcticus* (transcriptome) † | ERR2587155 | Brain |
|  | ERR2587156 | Heart |
|  | ERR2587157 | Kidney |
|  | ERR2587158 | Liver |
|  | ERR2587159 | Skin |
|  | ERR2587160 | White Muscle |
| *Parachaenichthys charcoti* (genome) ^x^ | doi:10.5524/100321 | N/A |
| *Parachaenichthys charcoti* (transcriptome)† | ERR2587183 | Blood |
|  | ERR2587164 | Brain |
|  | ERR2587166 | Head kidney |
|  | ERR2587169 | Liver |
|  | ERR2587171 | Ovary |
|  | ERR2587174 | Spleen |
|  | ERR2587177 | Trunk kidney |
|  | ERR2587179 | Ventricle |
|  | ERR2587181 | White Muscle |
| *Gymnodraco acuticeps* (transcriptome) | SRR6450839 | Pooled embryo and adult tissues |
|  | SRR6450835 | Pooled embryo and adult tissues |
|  | SRR6450836 | Pooled embryo tissues |
|  | SRR6450837 | Pooled embryo tissues |
|  | SRR6450838 | adult tissue pool (brain, gill, liver, spleen) |
| *Pseudochaenichthys georgianus* (transcriptome) † | ERR2587183 | Blood |
|  | ERR2587177 | Caudal Kidney |
|  | ERR2587166 | Head kidney |
|  | ERR2587179 | Heart (Ventricle) |
|  | ERR2587169 | Liver |
|  | ERR2587172 | Muscle (Pectoral) |
|  | ERR2587181 | Muscle (White) |
|  | ERR2587171 | Ovary |
|  | ERR2587174 | Spleen |
| *Chaenocephalus aceratus* (transcriptome) | SRR6929344 | Brain |
|  | SRR6929343 | Eye |
|  | SRR6929342 | Gill |
|  | SRR6929341 | Heart |
|  | SRR6929348 | Intestine |
|  | SRR6929347 | Kidney |
|  | SRR6929346 | Liver |
|  | SRR6929345 | Muscle |
|  | SRR6929350 | Ovary |
|  | SRR6929349 | Skin |
|  | SRR6929352 | Spleen |
|  | SRR6929351 | Stomach |
| *Chionodraco myersi* (transcriptome) | SRR8197049 | Liver |
|  | SRR8197051 | Brain |
|  | SRR8197052 | Kidney |
|  | SRR8197050 | Spleen |
| *Chionodraco hamatus* (transcriptome) | SRR4279902 | Gills |
|  | SRR3114175 | Heart |
|  | SRR3114176 | Muscle |
|  | SRR3112442 | Liver |
|  | SRR2072638 | N/A |
|  | SRR2072637 | N/A |
|  | SRR2072636 | N/A |
| Chionodraco rastrospinosus (transcriptome) | SRR5210463  (pool) | Liver  Gill  Brain  Spleen  Heart  Head Kidney  Stomach  Intestine  Eye Cup  Lens |

Unless otherwise noted sequences were downloaded from NCBI’s SRA.

Species denoted with † were downloaded from ArrayExpress E-MTAB-6759.

Species denoted with ^x^ were downloaded from GIGADB

Species denoted with ^|^ were downloaded from NCBI’s genome assembly database

**S Material 2. Genomic & Transcriptome sequences used in the data analysis (Page 1 of 2)**

| **Species** | **16S** | **ND2** | **HECW2** | **MYH6** | **PKD1** | **PPM1D** |
| --- | --- | --- | --- | --- | --- | --- |
| *Seriola dumerili* | NC_016870.1 | NC_016870.1 | ENSSDUP00000025534 ^1^ | ENSSDUT00000022504.1 ^1^ | ENSSDUT00000003940.1 ^1^ | N/A |
| *Labrus bergylta* | MT410912.1 | MT410912.1 | ENSLBEP00000029024 ^1^ | ENSLCRP00005003428 ^1^ | ENSLBET00000015305.1 ^1^ | N/A |
| *Takifugu rubripes* | AP006045.1 | AP006045.1 | ENSTRUP00000023073 ^1^ | ENSTRUP00000086471 ^1^ | N/A | N/A |
| *Larimichthys crocea* | NC_011710.1 | NC_011710.1 | ENSLCRP00005008096 ^1^ | ENSLCRP00005003428 ^1^ | ENSLCRT00005050827.1 ^1^ | N/A |
| *Gasterosteus aculeatus* | NC_041244.1 | NC_041244.1 | ENSGACP00000020039 ^1^ | ENSGACP00000018205 ^1^ | ENSGACT00000025106.1 ^1^ | N/A |
| *Cottoperca gobio* | AY520102.1 | JN186884.1 | JQ688794.1 | JN187021.1 | JQ688745.1 | N/A |
| *Eleginops maclovinus* | JN186915.1 | KF412874.1 | JQ688795.1 | JN187023.1 | JQ688746.1 | JQ688691.1 |
| *Dissostichus mawsoni* | LC138011.1 | AY256561.1 | JQ688816.1 | JN187040.1 | JQ688764.1 | JQ688715.1 |
| *Notothenia coriiceps* | NC_015653.1 | AY256563.1 | JQ688821.1 | JN187047.1 | JQ688757.1 | JQ688719.1 |
| *Pagothenia borchgrevinki* | KX025131.1 | KR153352.1 | JQ688817.1 | HM166364.1 | JQ688765.1 | † |
| *Trematomus bernacchii* | AY520126.1 | KR153464.1 | JQ688841.1 | HM166366.1 | JQ688784.1 | † |
| *Harpagifer antarcticus* | AY520130.1 | JN186894.1 | N/A | HQ169791.1 | † | JQ688717.1 |
| *Gymnodraco acuticeps* | MT559888.1 | MT559888.1 | JQ688811.1 | HQ169837.1 | JQ688759.1 | JQ688709.1 |
| *Parachaenichthys charcoti* | KP300644.1 | KP300644.1 | N/A | HQ169840.1 | † | N/A |
| *Chaenocephalus aceratus* | F933907.1 | JF933907.1 | JQ688805.1 | HQ169793.1 | † | JQ688701.1 |
| *Pseudochaenichthys georgianus* | MT559893.1 | MT559893.1 | N/A | JF264547.1 | † | N/A |
| *Chionodraco hamatus* | HQ170305.1 | HQ170101.1 | JQ688815.1 | HQ169804.1 | JQ688763.1 | † |
| *Chionodraco rastrospinosus* | AY249475.1 | AY249502.1 | N/A | HQ169808.1 | † | N/A |
| *Chionodraco myersi* | DQ526430.1 | HQ170103.1 | N/A | HQ169807.1 | † | N/A |

**S Material 2. Genomic & Transcriptome sequences used in the data analysis (Page 2 of 2)**

| **Species** | **PTR** | **RHO** | **SH3PX3** | **SSRP1** | **TBR1** | **ZIC1** | **RPS71** |
| --- | --- | --- | --- | --- | --- | --- | --- |
| *Seriola dumerili* | ENSSDUT00000012076.1 ^1^ | ENSSDUT00000019767.1 ^1^ | XM_022756080.1 | ENSSDUT00000020584.1 ^1^ | XM_022740861.1 | XM_022767191.1 | N/A |
| *Labrus bergylta* | ENSLBET00000037972.1 ^1^ | ENSLBET00000012479.1 ^1^ | XM_020660907.2 | ENSLBET00000013542.1 ^1^ | XM_020644160.2 | XM_020654969.2 | N/A |
| *Takifugu rubripes* | ENSTRUT00000044496.3 ^1^ | ENSTRUT00000010811.3 ^1^ | XM_003967599.3 | ENSTRUT00000033688.3 ^1^ | XM_003961934.2 | XM_003968020.3 | N/A |
| *Larimichthys crocea* | ENSLCRT00005025024.1 ^1^ | ENSLCRT00005061767.1 ^1^ | XM_010735182.3 | ENSLCRT00005017928.1 ^1^ | XM_010732216.3 | XM_010753423.3 | N/A |
| *Gasterosteus aculeatus* | ENSGACT00000005605.1 ^1^ | EU637962.1 | XM_040162897.1 | ENSGACT00000023573.1 ^1^ | XM_040200671.1 | XM_040167334.1 | N/A |
| *Cottoperca gobio* | ENSCGOT00000017205.1 ^1^ | ENSCGOT00000016480.1 ^1^ | JN186939.1 | N/A | XM_029459088.1 | JN186803.1 | JN186844.1 |
| *Eleginops maclovinus* | HM050163.1 | AY141303.1 | JN186941.1 | N/A | JN186982.1 | JN186805.1 | JN186846.1 |
| *Dissostichus mawsoni* | JF264567.1 | DQ498794.1 | JN186958.1 | N/A | JN186999.1 | JN186822.1 | AY517753.1 |
| *Notothenia coriiceps* | HM050183.1 | XM_010795871.1 | JN186965.1 | N/A | JN187006.1 | XM_010793969.1 | FJ647673.1 |
| *Pagothenia borchgrevinki* | HM166315.1 | GU997240.1 | HM166239.1 | N/A | HM166214.1 | HM166190.1 | FJ647674.1 |
| *Trematomus bernacchii* | HM166317.1 | DQ498797.1 | HM166241.1 | N/A | HM050267.1 | HM166191.1 | FJ647676.1 |
| *Harpagifer antarcticus* | HQ169911.1 | HQ170033.1 | HQ170178.1 | N/A | HQ170237.1 | HQ169671.1 | HQ170134.1 |
| *Gymnodraco acuticeps* | HQ169958.1 | KU647485.1 | XM_034236151.1 | N/A | JF264606.1 | HQ169718.1 | HQ170165.1 |
| *Parachaenichthys charcoti* | HQ169960.1 | HQ170081.1 | HQ170226.1 | N/A | KC830426.1 | HQ169720.1 | HQ170167.1 |
| *Chaenocephalus aceratus* | HQ169914.1 | HQ170036.1 | HQ170180.1 | N/A | HQ170239.1 | HQ169674.1 | HM166091.1 |
| *Pseudochaenichthys georgianus* | HQ169943.1 | HQ170065.1 | HQ170210.1 | N/A | JF264622.1 | HQ169703.1 | HM165578.1 |
| *Chionodraco hamatus* | HQ169925.1 | HQ170047.1 | HQ170191.1 | N/A | HQ170250.1 | HQ169684.1 | HQ170142.1 |
| *Chionodraco rastrospinosus* | HQ169929.1 | HQ170050.1 | HQ170196.1 | N/A | KC830288.1 | HQ169688.1 | HM165855.1 |
| *Chionodraco myersi* | HQ169927.1 | HQ170048.1 | HQ170194.1 | N/A | HQ170049.1 | HQ169687.1 | HQ170144.1 |

^1^ denotes genes from ensemble, all other accession numbers reference NCBI’s genbank. Genes denoted with † were isolated from draft genomic annotations or transcriptomic assemblies. N/A is used to denote species gene combinations that were not included in the analysis.

**S Material 3. Phylogenetic reconstruction of the species used in this study**

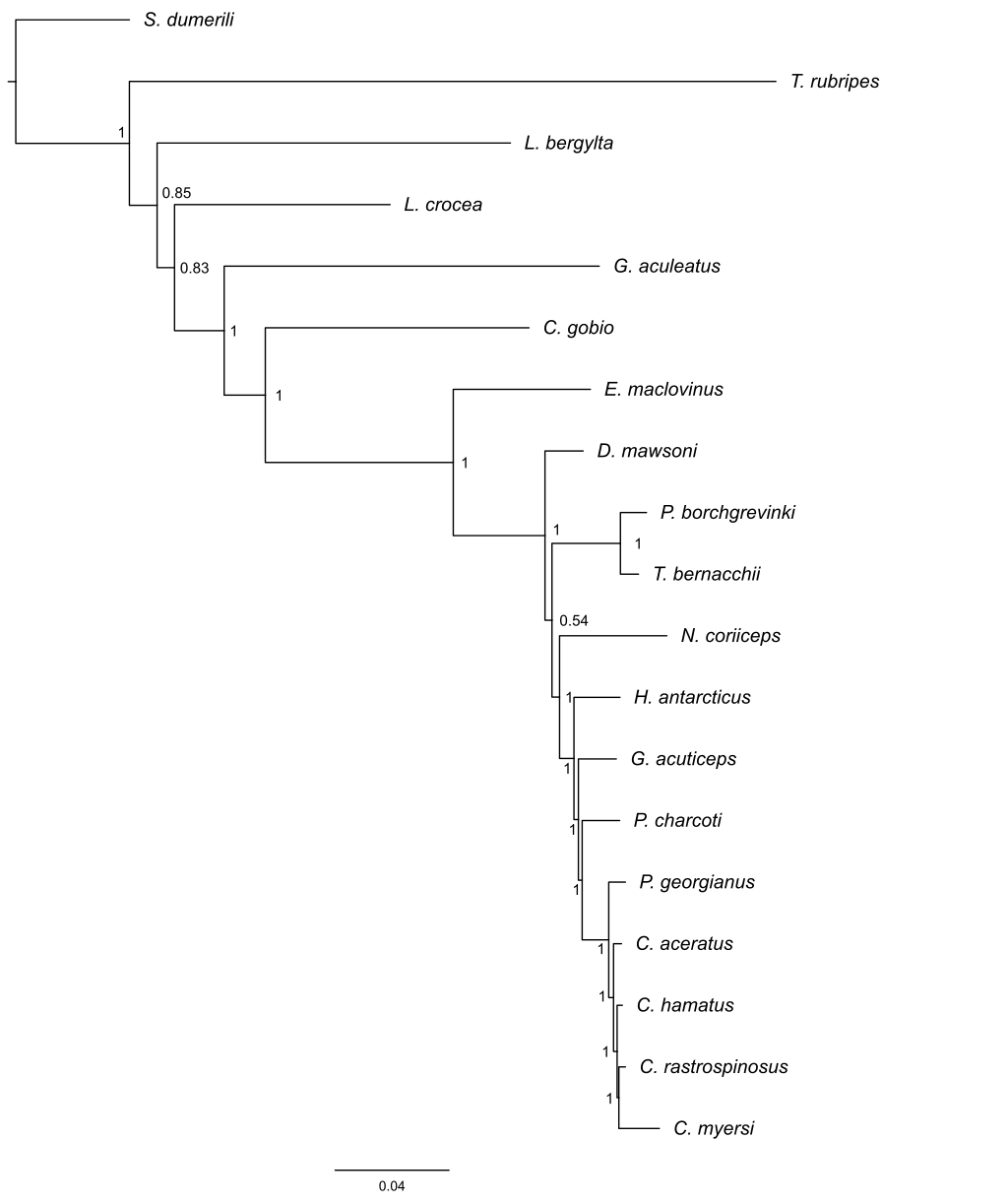

**S Material 4. Tissue completeness of each notothenioid species’ transcriptome**

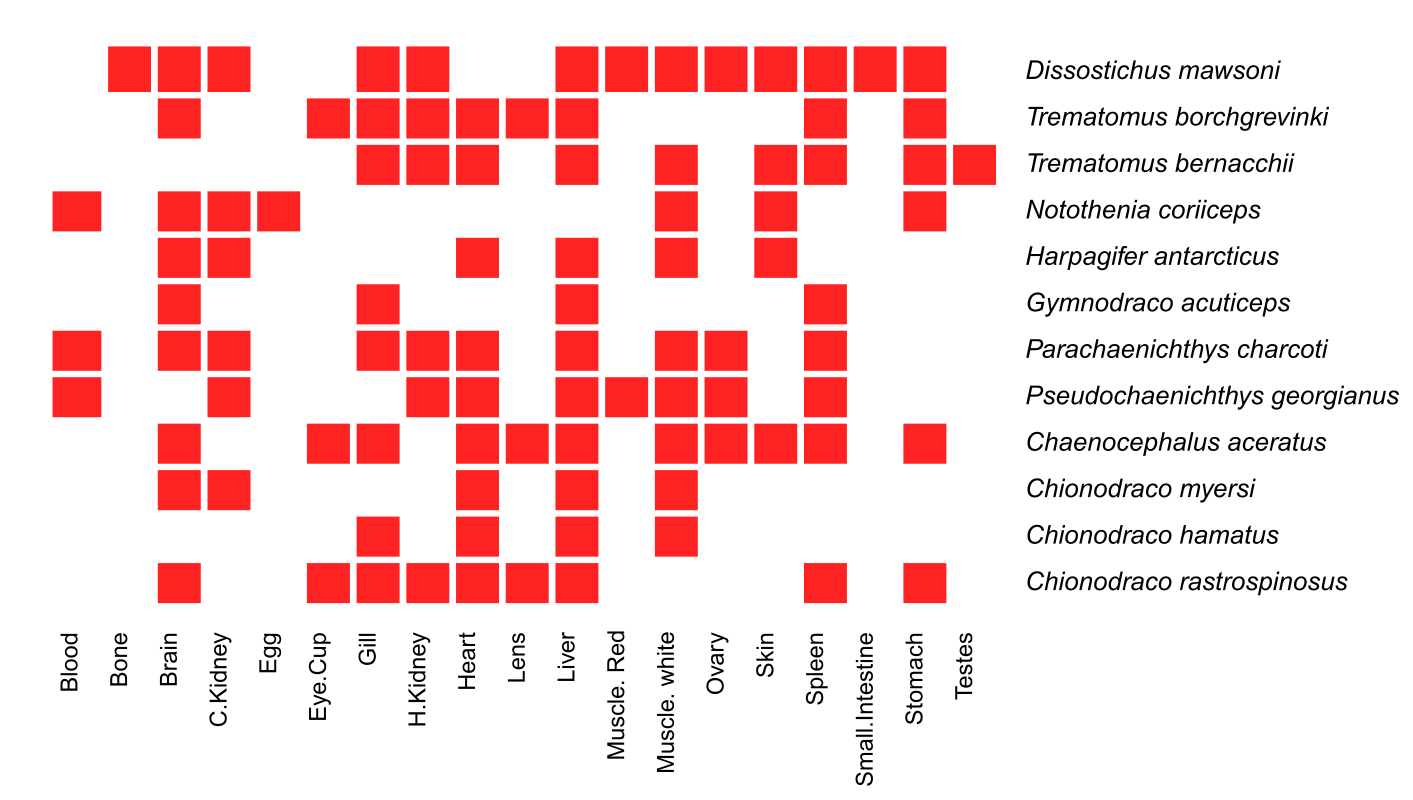

**S Material 5. Comparison of genomic and transcriptomic derived predicted peptides using BUSCO for three cryonotothenioid species**

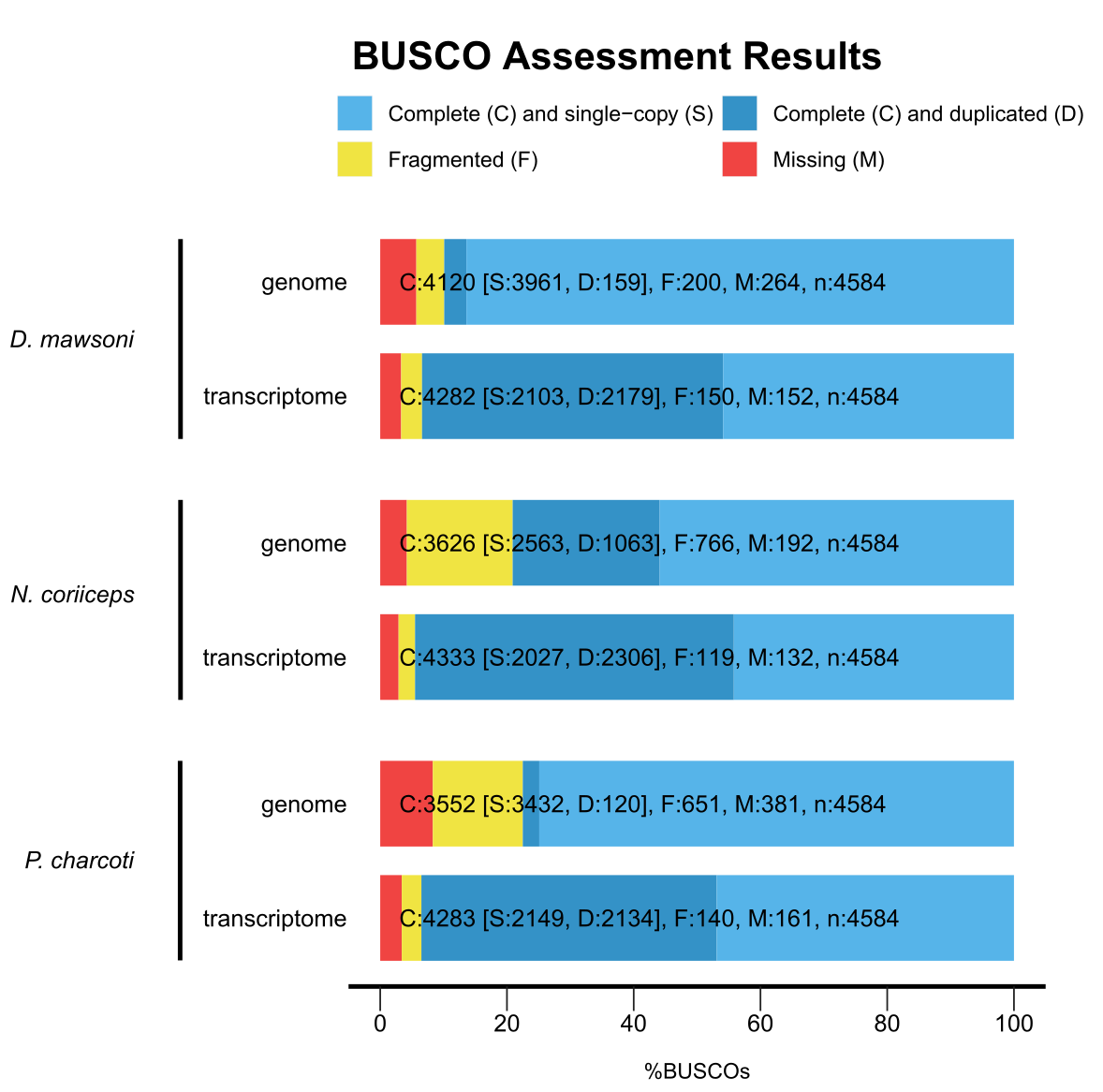

**S Material 6:**

**Enriched GO terms for orthogroups identified under positive diversifying selective pressure by either (union) BUSTED or aBSREL in the red blooded cryonotothenioids**

**Biological Process (BP)**

| **GO** | **term** | **Annotated** | **Significant** | **P-value** |
| --- | --- | --- | --- | --- |
| GO:0006355 | regulation of transcription, DNA-templat… | 371 | 29 | 0.0019 |
| GO:1903047 | mitotic cell cycle process | 93 | 10 | 0.0088 |
| GO:0098609 | cell-cell adhesion | 42 | 6 | 0.0109 |
| GO:0009408 | response to heat | 12 | 3 | 0.0151 |
| GO:0140014 | mitotic nuclear division | 34 | 5 | 0.0175 |
| GO:0022610 | biological adhesion | 89 | 9 | 0.0185 |
| GO:0007155 | cell adhesion | 89 | 9 | 0.0185 |
| GO:0014070 | response to organic cyclic compound | 36 | 5 | 0.0221 |
| GO:2001056 | positive regulation of cysteine-type end... | 14 | 3 | 0.0233 |
| GO:0010950 | positive regulation of endopeptidase act... | 14 | 3 | 0.0233 |
| GO:0010952 | positive regulation of peptidase activit... | 14 | 3 | 0.0233 |
| GO:0007052 | mitotic spindle organization | 14 | 3 | 0.0233 |
| GO:0043280 | positive regulation of cysteine-type end... | 14 | 3 | 0.0233 |
| GO:0032271 | regulation of protein polymerization | 14 | 3 | 0.0233 |
| GO:0000280 | nuclear division | 38 | 5 | 0.0273 |
| GO:0008015 | blood circulation | 26 | 4 | 0.0283 |
| GO:0042157 | lipoprotein metabolic process | 26 | 4 | 0.0283 |
| GO:2000113 | negative regulation of cellular macromol... | 130 | 11 | 0.0325 |
| GO:0048285 | organelle fission | 40 | 5 | 0.0333 |
| GO:0051258 | protein polymerization | 16 | 3 | 0.0336 |
| GO:0003013 | circulatory system process | 28 | 4 | 0.0362 |
| GO:0010558 | negative regulation of macromolecule bio... | 133 | 11 | 0.0376 |
| GO:0045862 | positive regulation of proteolysis | 17 | 3 | 0.0395 |
| GO:1902903 | regulation of supramolecular fiber organ... | 17 | 3 | 0.0395 |
| GO:0051092 | positive regulation of NF-kappaB transcr... | 17 | 3 | 0.0395 |
| GO:0031327 | negative regulation of cellular biosynth... | 136 | 11 | 0.0432 |
| GO:0007346 | regulation of mitotic cell cycle | 72 | 7 | 0.0436 |
| GO:0009890 | negative regulation of biosynthetic proc... | 137 | 11 | 0.0452 |
| GO:0009266 | response to temperature stimulus | 18 | 3 | 0.0458 |
| GO:0043254 | regulation of protein-containing complex... | 18 | 3 | 0.0458 |

**Molecular Function (MF)**

| **GO** | **term** | **Annotated** | **Significant** | **P-value** |
| --- | --- | --- | --- | --- |
| GO:0050839 | cell adhesion molecule binding | 23 | 4 | 0.019 |
| GO:0016651 | oxidoreductase activity, acting on NAD(P... | 13 | 3 | 0.019 |
| GO:0001664 | G protein-coupled receptor binding | 14 | 3 | 0.023 |
| GO:0003700 | DNA-binding transcription factor activit... | 116 | 10 | 0.036 |

**Cellular Component (CC)**

| **GO** | **term** | **Annotated** | **Significant** | **P-value** |
| --- | --- | --- | --- | --- |
| GO:0019897 | extrinsic component of plasma membrane | 20 | 5 | 0.0015 |
| GO:0005911 | cell-cell junction | 30 | 5 | 0.0096 |

**S Material 7:**

**Enriched GO terms for orthogroups identified under positive diversifying selective pressure by only BUSTED in the red blooded cryonotothenioids**

**Biological Process (BP)**

| **GO** | **term** | **Annotated** | **Significant** | **P-value** |
| --- | --- | --- | --- | --- |
| GO:0042157 | lipoprotein metabolic process | 26 | 4 | 0.0030 |
| GO:0014070 | response to organic cyclic compound | 36 | 4 | 0.0099 |
| GO:0042391 | regulation of membrane potential | 20 | 3 | 0.0111 |
| GO:0050727 | regulation of inflammatory response | 22 | 3 | 0.0145 |
| GO:0042493 | response to drug | 10 | 2 | 0.0224 |
| GO:0071901 | negative regulation of protein serine/th... | 10 | 2 | 0.0224 |
| GO:1901607 | alpha-amino acid biosynthetic process | 11 | 2 | 0.0269 |
| GO:0032465 | regulation of cytokinesis | 11 | 2 | 0.0269 |
| GO:0007187 | G protein-coupled receptor signaling pat... | 11 | 2 | 0.0269 |
| GO:0007188 | adenylate cyclase-modulating G protein-c... | 11 | 2 | 0.0269 |
| GO:1903707 | negative regulation of hemopoiesis | 11 | 2 | 0.0269 |
| GO:0007346 | regulation of mitotic cell cycle | 72 | 5 | 0.0273 |
| GO:0031348 | negative regulation of defense response | 12 | 2 | 0.0318 |
| GO:0071222 | cellular response to lipopolysaccharide | 12 | 2 | 0.0318 |
| GO:0071216 | cellular response to biotic stimulus | 12 | 2 | 0.0318 |
| GO:0071219 | cellular response to molecule of bacteri... | 12 | 2 | 0.0318 |
| GO:0008652 | cellular amino acid biosynthetic process | 12 | 2 | 0.0318 |
| GO:0009617 | response to bacterium | 31 | 3 | 0.0364 |
| GO:0071407 | cellular response to organic cyclic comp... | 31 | 3 | 0.0364 |
| GO:0030832 | regulation of actin filament length | 13 | 2 | 0.0370 |
| GO:0030833 | regulation of actin filament polymerizat... | 13 | 2 | 0.0370 |
| GO:0008064 | regulation of actin polymerization or de... | 13 | 2 | 0.0370 |
| GO:0008154 | actin polymerization or depolymerization | 13 | 2 | 0.0370 |
| GO:0030041 | actin filament polymerization | 13 | 2 | 0.0370 |
| GO:0043405 | regulation of MAP kinase activity | 13 | 2 | 0.0370 |
| GO:0033673 | negative regulation of kinase activity | 32 | 3 | 0.0395 |
| GO:0006469 | negative regulation of protein kinase ac... | 32 | 3 | 0.0395 |
| GO:0033043 | regulation of organelle organization | 107 | 6 | 0.0405 |
| GO:0060047 | heart contraction | 14 | 2 | 0.0425 |
| GO:0032271 | regulation of protein polymerization | 14 | 2 | 0.0425 |
| GO:0051348 | negative regulation of transferase activ... | 33 | 3 | 0.0427 |
| GO:1901701 | cellular response to oxygen-containing c... | 33 | 3 | 0.0427 |
| GO:0045930 | negative regulation of mitotic cell cycl... | 33 | 3 | 0.0427 |
| GO:0070887 | cellular response to chemical stimulus | 167 | 8 | 0.0430 |
| GO:0045892 | negative regulation of transcription, DN... | 110 | 6 | 0.0454 |
| GO:0051726 | regulation of cell cycle | 139 | 7 | 0.0457 |
| GO:0140014 | mitotic nuclear division | 34 | 3 | 0.0460 |
| GO:0043409 | negative regulation of MAPK cascade | 15 | 2 | 0.0483 |

**Molecular Function (MF)**

| **GO** | **Term** | **Annotated** | **Significant** | **P-value** |
| --- | --- | --- | --- | --- |
| GO:0016787 | hydrolase activity | 507 | 19 | 0.031 |
| GO:0072509 | divalent inorganic cation transmembrane ... | 12 | 2 | 0.033 |
| GO:0001664 | G protein-coupled receptor binding | 14 | 2 | 0.045 |
| GO:0003743 | translation initiation factor activity | 14 | 2 | 0.045 |
| GO:0008138 | protein tyrosine/serine/threonine phosph... | 14 | 2 | 0.045 |
| GO:0008237 | metallopeptidase activity | 33 | 3 | 0.046 |

**Cellular Component (CC)**

| **GO** | **Term** | **Annotated** | **Significant** | **P-value** |
| --- | --- | --- | --- | --- |
| GO:0072686 | mitotic spindle | 15 | 2 | 0.046 |

**S Material 8:**

**Enriched GO terms for orthogroups identified under positive diversifying selective pressure by only aBSREL in the red blooded cryonotothenioids**

**Biological Process (BP)**

| **GO** | **term** | **Annotated** | **Significant** | **P-value** |
| --- | --- | --- | --- | --- |
| GO:0006355 | regulation of transcription, DNA-templat... | 371 | 25 | 0.00023 |
| GO:2000113 | negative regulation of cellular macromol... | 130 | 10 | 0.00934 |
| GO:0042157 | lipoprotein metabolic process | 26 | 4 | 0.00946 |
| GO:0043280 | positive regulation of cysteine-type end... | 14 | 3 | 0.00968 |
| GO:0098609 | cell-cell adhesion | 42 | 5 | 0.01129 |
| GO:0051094 | positive regulation of developmental pro... | 64 | 6 | 0.01748 |
| GO:0051172 | negative regulation of nitrogen compound... | 188 | 12 | 0.01871 |
| GO:0022610 | biological adhesion | 89 | 7 | 0.02560 |
| GO:0007155 | cell adhesion | 89 | 7 | 0.02560 |
| GO:0031503 | protein-containing complex localization | 20 | 3 | 0.02628 |
| GO:0045892 | negative regulation of transcription, DN... | 110 | 8 | 0.02660 |
| GO:0051240 | positive regulation of multicellular org... | 72 | 6 | 0.02955 |
| GO:1903507 | negative regulation of nucleic acid-temp... | 113 | 8 | 0.03066 |
| GO:1902679 | negative regulation of RNA biosynthetic ... | 113 | 8 | 0.03066 |
| GO:0051495 | positive regulation of cytoskeleton orga... | 10 | 2 | 0.04068 |
| GO:0038061 | NIK/NF-kappaB signaling | 10 | 2 | 0.04068 |
| GO:0010257 | NADH dehydrogenase complex assembly | 10 | 2 | 0.04068 |
| GO:0018198 | peptidyl-cysteine modification | 10 | 2 | 0.04068 |
| GO:0071901 | negative regulation of protein serine/th... | 10 | 2 | 0.04068 |
| GO:1901222 | regulation of NIK/NF-kappaB signaling | 10 | 2 | 0.04068 |
| GO:1902905 | positive regulation of supramolecular fi... | 10 | 2 | 0.04068 |
| GO:0003018 | vascular process in circulatory system | 10 | 2 | 0.04068 |
| GO:0032981 | mitochondrial respiratory chain complex ... | 10 | 2 | 0.04068 |
| GO:0051253 | negative regulation of RNA metabolic pro... | 122 | 8 | 0.04537 |
| GO:0032269 | negative regulation of cellular protein ... | 80 | 6 | 0.04614 |
| GO:0051248 | negative regulation of protein metabolic... | 80 | 6 | 0.04614 |
| GO:0006457 | protein folding | 25 | 3 | 0.04867 |
| GO:0006919 | activation of cysteine-type endopeptidas... | 11 | 2 | 0.04867 |
| GO:0045934 | negative regulation of nucleobase-contai... | 124 | 8 | 0.04918 |

**Molecular Function (MF)**

| **GO** | **Term** | **Annotated** | **Significant** | **P-value** |
| --- | --- | --- | --- | --- |
| GO:0050839 | cell adhesion molecule binding | 23 | 4 | 0.0063 |
| GO:0016651 | oxidoreductase activity, acting on NAD(P... | 13 | 3 | 0.0080 |
| GO:0003700 | DNA-binding transcription factor activit... | 116 | 9 | 0.0137 |
| GO:0008173 | RNA methyltransferase activity | 19 | 3 | 0.0236 |
| GO:0003677 | DNA binding | 236 | 13 | 0.0462 |
| GO:0031406 | carboxylic acid binding | 25 | 3 | 0.0485 |
| GO:0043177 | organic acid binding | 25 | 3 | 0.0485 |
| GO:0002020 | protease binding | 11 | 2 | 0.0497 |

**Cellular Component (CC)**

| **GO** | **Term** | **Annotated** | **Significant** | **P-value** |
| --- | --- | --- | --- | --- |
| GO:0005911 | cell-cell junction | 30 | 4 | 0.016 |
| GO:0019897 | extrinsic component of plasma membrane | 20 | 3 | 0.027 |

**S Material 9:**

**Enriched GO terms for orthogroups consistently identified under positive diversifying selective pressure by both (intersection) BUSTED and aBSREL in the red blooded cryonotothenioids**

**Biological Process (BP)**

| **GO** | **term** | **Annotated** | **Significant** | **P-value** |
| --- | --- | --- | --- | --- |
| GO:0042157 | lipoprotein metabolic process | 26 | 4 | 0.00016 |
| GO:0071901 | negative regulation of protein serine/th... | 10 | 2 | 0.00519 |
| GO:0045892 | negative regulation of transcription, DN... | 110 | 5 | 0.00679 |
| GO:0043405 | regulation of MAP kinase activity | 13 | 2 | 0.00881 |
| GO:0043409 | negative regulation of MAPK cascade | 15 | 2 | 0.01169 |
| GO:0006497 | protein lipidation | 19 | 2 | 0.01852 |
| GO:0042158 | lipoprotein biosynthetic process | 20 | 2 | 0.02043 |
| GO:0051169 | nuclear transport | 30 | 2 | 0.04364 |
| GO:0006913 | nucleocytoplasmic transport | 30 | 2 | 0.04364 |

**Molecular Function (MF)**

| **GO** | **Term** | **Annotated** | **Significant** | **P-value** |
| --- | --- | --- | --- | --- |
| GO:0016747 | transferase activity, transferring acyl ... | 46 | 3 | 0.018 |
| GO:0016810 | hydrolase activity, acting on carbon-nit... | 22 | 2 | 0.029 |
| GO:0016746 | transferase activity, transferring acyl ... | 56 | 3 | 0.030 |
| GO:0016758 | transferase activity, transferring hexos... | 25 | 2 | 0.037 |
| GO:0043177 | organic acid binding | 25 | 2 | 0.037 |
| GO:0031406 | carboxylic acid binding | 25 | 2 | 0.037 |
| GO:0003690 | double-stranded DNA binding | 113 | 4 | 0.047 |
| GO:0016407 | acetyltransferase activity | 29 | 2 | 0.048 |

**Cellular Component (CC)**

None

**S Material 10:**

**Enriched GO terms for orthogroups consistently identified under relaxed selective pressure by RELAX in the red blooded cryonotothenioids**

**Biological Process (BP)**

| **GO** | **term** | **Annotated** | **Significant** | **P-value** |
| --- | --- | --- | --- | --- |
| GO:0060042 | retina morphogenesis in camera-type eye | 17 | 5 | 0.0013 |
| GO:0015711 | organic anion transport | 46 | 8 | 0.0022 |
| GO:0022904 | respiratory electron transport chain | 13 | 4 | 0.0035 |
| GO:0006476 | protein deacetylation | 14 | 4 | 0.0047 |
| GO:0048703 | embryonic viscerocranium morphogenesis | 15 | 4 | 0.0062 |
| GO:0006955 | Immune response | 85 | 10 | 0.0117 |
| GO:0003407 | neural retina development | 10 | 3 | 0.0127 |
| GO:0046942 | carboxylic acid transport | 10 | 3 | 0.0127 |
| GO:0090288 | negative regulation of cellular response... | 10 | 3 | 0.0127 |
| GO:0015849 | organic acid transport | 10 | 3 | 0.0127 |
| GO:0032446 | protein modification by small protein co... | 114 | 12 | 0.0141 |
| GO:0015748 | organophosphate ester transport | 19 | 4 | 0.0149 |
| GO:0006302 | double-strand break repair | 19 | 4 | 0.0149 |
| GO:0070647 | protein modification by small protein co... | 143 | 14 | 0.0151 |
| GO:0090101 | negative regulation of transmembrane rec... | 11 | 3 | 0.0168 |
| GO:0007409 | axonogenesis | 30 | 5 | 0.0178 |
| GO:0050776 | regulation of immune response | 41 | 6 | 0.0179 |
| GO:0006281 | DNA repair | 66 | 8 | 0.0198 |
| GO:0000003 | reproduction | 66 | 8 | 0.0198 |
| GO:0016567 | protein ubiquitination | 107 | 11 | 0.0217 |
| GO:0006974 | cellular response to DNA damage stimulus | 109 | 11 | 0.0245 |
| GO:0006259 | DNA metabolic process | 82 | 9 | 0.0250 |
| GO:0071772 | response to BMP | 23 | 4 | 0.0289 |
| GO:0071773 | cellular response to BMP stimulus | 23 | 4 | 0.0289 |
| GO:0030509 | BMP signaling pathway | 23 | 4 | 0.0289 |
| GO:0065008 | regulation of biological quality | 329 | 25 | 0.0305 |
| GO:0061564 | axon development | 36 | 5 | 0.0367 |
| GO:0022414 | reproductive process | 49 | 6 | 0.0397 |
| GO:0030510 | regulation of BMP signaling pathway | 15 | 3 | 0.0398 |
| GO:0048667 | cell morphogenesis involved in neuron di... | 37 | 5 | 0.0407 |
| GO:0051216 | cartilage development | 37 | 5 | 0.0407 |
| GO:0000904 | cell morphogenesis involved in different... | 50 | 6 | 0.0433 |
| GO:0006869 | lipid transport | 26 | 4 | 0.0433 |
| GO:0009790 | embryo development | 169 | 17 | 0.0469 |
| GO:0007519 | skeletal muscle tissue development | 16 | 3 | 0.0472 |
| GO:0071478 | cellular response to radiation | 16 | 3 | 0.0472 |
| GO:0061448 | connective tissue development | 39 | 5 | 0.0495 |
| GO:0120039 | plasma membrane bounded cell projection ... | 39 | 5 | 0.0495 |
| GO:0048812 | neuron projection morphogenesis | 39 | 5 | 0.0495 |

**Molecular Function (MF)**

| **GO** | **term** | **Annotated** | **Significant** | **P-value** |
| --- | --- | --- | --- | --- |
| GO:0003677 | DNA binding | 236 | 21 | 0.0083 |
| GO:0000981 | DNA-binding transcription factor activit... | 72 | 9 | 0.0107 |
| GO:0003690 | double-stranded DNA binding | 113 | 12 | 0.0123 |
| GO:0003684 | damaged DNA binding | 11 | 3 | 0.0165 |
| GO:0005215 | transporter activity | 152 | 14 | 0.0230 |
| GO:0015078 | proton transmembrane transporter activit... | 22 | 4 | 0.0242 |
| GO:0015075 | ion transmembrane transporter activity | 100 | 10 | 0.0316 |
| GO:0001228 | DNA-binding transcription activator acti... | 8 | 24 | 0.0325 |
| GO:0001216 | DNA-binding transcription activator acti... | 24 | 4 | 0.0325 |
| GO:0001067 | regulatory region nucleic acid binding | 87 | 9 | 0.0335 |
| GO:0000976 | transcription regulatory region sequence... | 87 | 9 | 0.0335 |
| GO:0003700 | DNA-binding transcription factor activit... | 116 | 11 | 0.0349 |
| GO:1990837 | sequence-specific double-stranded DNA bi... | 88 | 9 | 0.0358 |
| GO:0008514 | organic anion transmembrane transporter ... | 38 | 5 | 0.0437 |
| GO:0003697 | single-stranded DNA binding | 16 | 3 | 0.0463 |

**Cellular Component (CC)**

| **GO** | **term** | **Annotated** | **Significant** | **P-value** |
| --- | --- | --- | --- | --- |
| GO:0005759 | mitochondrial matrix | 80 | 13 | 0.00032 |
| GO:0005746 | mitochondrial respirasome | 12 | 4 | 0.00309 |

**S Material 11:**

**Enriched GO terms for orthogroups identified under positive diversifying selective pressure by either (union) BUSTED or aBSREL in the icefishes**

**Biological Process (BP)**

| **GO** | **term** | **Annotated** | **Significant** | **P-value** |
| --- | --- | --- | --- | --- |
| GO:0019725 | cellular homeostasis | 96 | 5 | 0.0050 |
| GO:1901222 | regulation of NIK/NF-kappaB signaling | 10 | 2 | 0.0059 |
| GO:0007188 | adenylate cyclase-modulating G protein-c... | 11 | 2 | 0.0071 |
| GO:0003143 | embryonic heart tube morphogenesis | 12 | 2 | 0.0085 |
| GO:0006006 | glucose metabolic process | 16 | 2 | 0.0150 |
| GO:0016051 | carbohydrate biosynthetic process | 17 | 2 | 0.0168 |
| GO:0033500 | carbohydrate homeostasis | 19 | 2 | 0.0209 |
| GO:0042593 | glucose homeostasis | 19 | 2 | 0.0209 |
| GO:0019318 | hexose metabolic process | 21 | 2 | 0.0252 |
| GO:0005996 | monosaccharide metabolic process | 21 | 2 | 0.0252 |
| GO:0008213 | protein alkylation | 22 | 2 | 0.0276 |
| GO:0048871 | multicellular organismal homeostasis | 22 | 2 | 0.0276 |
| GO:0006479 | protein methylation | 22 | 2 | 0.0276 |
| GO:0001837 | epithelial to mesenchymal transition | 22 | 2 | 0.0276 |
| GO:0009719 | response to endogenous stimulus | 98 | 4 | 0.0277 |
| GO:0009725 | response to hormone | 57 | 3 | 0.0293 |
| GO:0006898 | receptor-mediated endocytosis | 23 | 2 | 0.0300 |
| GO:0051093 | negative regulation of developmental pro... | 60 | 3 | 0.0335 |
| GO:0006869 | lipid transport | 26 | 2 | 0.0376 |
| GO:0005975 | carbohydrate metabolic process | 63 | 3 | 0.0379 |
| GO:0051241 | negative regulation of multicellular org... | 65 | 3 | 0.0411 |
| GO:0060249 | anatomical structure homeostasis | 28 | 2 | 0.0431 |
| GO:0010876 | lipid localization | 28 | 2 | 0.0431 |
| GO:0060828 | regulation of canonical Wnt signaling pa... | 29 | 2 | 0.0460 |

**Molecular Function (MF)**

| **GO** | **Term** | **Annotated** | **Significant** | **P-value** |
| --- | --- | --- | --- | --- |
| GO:0008146 | sulfotransferase activity | 14 | 2 | 0.0092 |
| GO:0003924 | GTPase activity | 74 | 4 | 0.0093 |
| GO:0097367 | carbohydrate derivative binding | 395 | 9 | 0.0285 |
| GO:0060089 | molecular transducer activity | 62 | 3 | 0.0329 |
| GO:0038023 | signaling receptor activity | 62 | 3 | 0.0329 |
| GO:0000166 | nucleotide binding | 428 | 9 | 0.0454 |
| GO:1901265 | nucleoside phosphate binding | 428 | 9 | 0.0454 |

**Cellular Component (CC)**

| **GO** | **Term** | **Annotated** | **Significant** | **P-value** |
| --- | --- | --- | --- | --- |
| GO:0005886 | plasma membrane | 365 | 10 | 0.0056 |
| GO:0030424 | axon | 26 | 2 | 0.0348 |
| GO:0005911 | cell-cell junction | 30 | 2 | 0.0453 |

**S Material 12:**

**Enriched GO terms for orthogroups identified under positive diversifying selective pressure by only BUSTED in the icefishes**

**Biological Process (BP)**

| **GO** | **term** | **Annotated** | **Significant** | **P-value** |
| --- | --- | --- | --- | --- |
| GO:0006006 | glucose metabolic process | 16 | 2 | 0.0020 |
| GO:0006479 | protein methylation | 22 | 2 | 0.0038 |
| GO:0009966 | regulation of signal transduction | 308 | 5 | 0.0068 |
| GO:0019725 | cellular homeostasis | 96 | 3 | 0.0071 |
| GO:0018205 | peptidyl-lysine modification | 46 | 2 | 0.0161 |
| GO:0018193 | peptidyl-amino acid modification | 130 | 3 | 0.0164 |
| GO:0009725 | response to hormone | 57 | 2 | 0.0242 |
| GO:0030334 | regulation of cell migration | 59 | 2 | 0.0258 |
| GO:2000145 | regulation of cell motility | 65 | 2 | 0.0309 |
| GO:0040012 | regulation of locomotion | 65 | 2 | 0.0309 |
| GO:0051270 | regulation of cellular component movemen... | 67 | 2 | 0.0327 |
| GO:0006325 | chromatin organization | 68 | 2 | 0.0336 |
| GO:0048925 | lateral line system development | 10 | 1 | 0.0427 |
| GO:0010906 | regulation of glucose metabolic process | 10 | 1 | 0.0427 |
| GO:0038061 | NIK/NF-kappaB signaling | 10 | 1 | 0.0427 |
| GO:1901222 | regulation of NIK/NF-kappaB signaling | 10 | 1 | 0.0427 |
| GO:0018022 | peptidyl-lysine methylation | 10 | 1 | 0.0427 |
| GO:0010675 | regulation of cellular carbohydrate meta... | 11 | 1 | 0.0468 |
| GO:0007187 | G protein-coupled receptor signaling pat... | 11 | 1 | 0.0468 |
| GO:0007188 | adenylate cyclase-modulating G protein-c... | 11 | 1 | 0.0468 |

**Molecular Function (MF)**

| **GO** | **term** | **Annotated** | **Significant** | **P-value** |
| --- | --- | --- | --- | --- |
| GO:0008276 | protein methyltransferase activity | 10 | 1 | 0.045 |
| GO:0004180 | carboxypeptidase activity | 10 | 1 | 0.045 |

**Cellular Component (CC)**

| **GO** | **Term** | **Annotated** | **Significant** | **P-value** |
| --- | --- | --- | --- | --- |
| GO:0005576 | extracellular region | 164 | 3 | 0.040 |

**S Material 13:**

**Enriched GO terms for orthogroups identified under positive diversifying selective pressure by only aBSREL in the icefishes**

**Biological Process (BP)**

| **GO** | **term** | **Annotated** | **Significant** | **P-value** |
| --- | --- | --- | --- | --- |
| GO:1901222 | regulation of NIK/NF-kappaB signaling | 10 | 2 | 0.0039 |
| GO:0007188 | adenylate cyclase-modulating G protein-c... | 11 | 2 | 0.0048 |
| GO:0019725 | cellular homeostasis | 96 | 4 | 0.0130 |
| GO:0009719 | response to endogenous stimulus | 98 | 4 | 0.0140 |
| GO:0033500 | carbohydrate homeostasis | 19 | 2 | 0.0142 |
| GO:0042593 | glucose homeostasis | 19 | 2 | 0.0142 |
| GO:0035239 | tube morphogenesis | 104 | 4 | 0.0171 |
| GO:0009725 | response to hormone | 57 | 3 | 0.0176 |
| GO:0048871 | multicellular organismal homeostasis | 22 | 2 | 0.0188 |
| GO:0001837 | epithelial to mesenchymal transition | 22 | 2 | 0.0188 |
| GO:0051093 | negative regulation of developmental pro... | 60 | 3 | 0.0197 |
| GO:0006898 | receptor-mediated endocytosis | 23 | 2 | 0.0205 |
| GO:0051241 | negative regulation of multicellular org... | 65 | 3 | 0.0243 |
| GO:0006869 | lipid transport | 26 | 2 | 0.0259 |
| GO:0035295 | tube development | 121 | 4 | 0.0282 |
| GO:0042592 | homeostatic process | 183 | 5 | 0.0296 |
| GO:0010876 | lipid localization | 28 | 2 | 0.0298 |
| GO:0048646 | anatomical structure formation involved ... | 127 | 4 | 0.0329 |
| GO:0003007 | heart morphogenesis | 31 | 2 | 0.0360 |
| GO:0060562 | epithelial tube morphogenesis | 33 | 2 | 0.0403 |
| GO:0048762 | mesenchymal cell differentiation | 36 | 2 | 0.0473 |
| GO:0055085 | transmembrane transport | 143 | 4 | 0.0478 |

**Molecular Function (MF)**

| **GO** | **term** | **Annotated** | **Significant** | **P-value** |
| --- | --- | --- | --- | --- |
| GO:0003924 | GTPase activity | 74 | 4 | 0.0041 |
| GO:0097367 | carbohydrate derivative binding | 395 | 9 | 0.0060 |
| GO:0000166 | nucleotide binding | 428 | 9 | 0.0103 |
| GO:1901265 | nucleoside phosphate binding | 428 | 9 | 0.0103 |
| GO:0016782 | transferase activity, transferring sulfu... | 21 | 2 | 0.0153 |
| GO:0035639 | purine ribonucleoside triphosphate bindi... | 377 | 8 | 0.0156 |
| GO:0032555 | purine ribonucleotide binding | 380 | 8 | 0.0164 |
| GO:0032553 | ribonucleotide binding | 383 | 8 | 0.0171 |
| GO:0017076 | purine nucleotide binding | 385 | 8 | 0.0176 |
| GO:0060089 | molecular transducer activity | 62 | 3 | 0.0181 |
| GO:0038023 | signaling receptor activity | 62 | 3 | 0.0181 |
| GO:0036094 | small molecule binding | 476 | 9 | 0.0203 |
| GO:0016740 | transferase activity | 481 | 9 | 0.0217 |
| GO:0043168 | anion binding | 488 | 9 | 0.0237 |
| GO:0022857 | transmembrane transporter activity | 139 | 4 | 0.0357 |
| GO:0004674 | protein serine/threonine kinase activity | 81 | 3 | 0.0365 |
| GO:0015267 | channel activity | 37 | 2 | 0.0444 |
| GO:0022803 | passive transmembrane transporter activi... | 37 | 2 | 0.0444 |
| GO:0005215 | transporter activity | 152 | 4 | 0.0473 |

**Cellular Component (CC)**

| **GO** | **term** | **Annotated** | **Significant** | **P-value** |
| --- | --- | --- | --- | --- |
| GO:0005886 | plasma membrane | 365 | 9 | 0.0019 |
| GO:0030424 | axon | 26 | 2 | 0.0199 |
| GO:0031226 | intrinsic component of plasma membrane | 76 | 3 | 0.0252 |
| GO:0005911 | cell-cell junction | 30 | 2 | 0.0260 |
| GO:0030054 | Cell junction | 99 | 3 | 0.0495 |

**S Material 14:**

**Enriched GO terms for orthogroups consistently identified under relaxed selective pressure by RELAX in the icefishes**

**Biological Process (BP)**

| **GO** | **term** | **Annotated** | **Significant** | **P-value** |
| --- | --- | --- | --- | --- |
| GO:0007033 | vacuole organization | 29 | 2 | 0.0066 |
| GO:0015672 | monovalent inorganic cation transport | 42 | 2 | 0.0135 |
| GO:0055085 | transmembrane transport | 143 | 3 | 0.0212 |
| GO:0098662 | inorganic cation transmembrane transport | 58 | 2 | 0.0250 |
| GO:0098660 | inorganic ion transmembrane transport | 64 | 2 | 0.0300 |
| GO:0098655 | cation transmembrane transport | 67 | 2 | 0.0327 |
| GO:0016311 | dephosphorylation | 72 | 2 | 0.0374 |
| GO:1905037 | autophagosome organization | 10 | 1 | 0.0427 |
| GO:0070121 | Kupffer's vesicle development | 10 | 1 | 0.0427 |
| GO:0018198 | peptidyl-cysteine modification | 10 | 1 | 0.0427 |
| GO:0000045 | autophagosome assembly | 10 | 1 | 0.0427 |
| GO:0060491 | regulation of cell projection assembly | 11 | 1 | 0.0468 |
| GO:0010389 | regulation of G2/M transition of mitotic... | 11 | 1 | 0.0468 |

**Molecular Function (MF)**

| **GO** | **term** | **Annotated** | **Significant** | **P-value** |
| --- | --- | --- | --- | --- |
| GO:0016788 | hydrolase activity, acting on ester bond... | 158 | 4 | 0.0044 |
| GO:0015077 | monovalent inorganic cation transmembran... | 33 | 2 | 0.0095 |
| GO:0022803 | passive transmembrane transporter activi... | 37 | 2 | 0.0118 |
| GO:0015267 | channel activity | 37 | 2 | 0.0118 |
| GO:0015399 | primary active transmembrane transporter... | 10 | 1 | 0.0451 |
| GO:0016409 | palmitoyltransferase activity | 10 | 1 | 0.0451 |
| GO:0019706 | protein-cysteine S-palmitoyltransferase ... | 10 | 1 | 0.0451 |
| GO:0019707 | protein-cysteine S-acyltransferase activ... | 10 | 1 | 0.0451 |
| GO:0016417 | S-acyltransferase activity | 10 | 1 | 0.0451 |
| GO:0140098 | catalytic activity, acting on RNA | 77 | 2 | 0.0469 |
| GO:0016791 | phosphatase activity | 78 | 2 | 0.0480 |

**Cellular Component (CC)**

| **GO** | **term** | **Annotated** | **Significant** | **P-value** |
| --- | --- | --- | --- | --- |
| GO:0016021 | integral component of membrane | 646 | 7 | 0.030 |
| GO:0031224 | intrinsic component of membrane | 650 | 7 | 0.031 |
